# Face dissimilarity judgements are predicted by representational distance in morphable and image-computable models

**DOI:** 10.1101/2021.04.09.438859

**Authors:** Kamila M. Jozwik, Jonathan O’Keeffe, Katherine R. Storrs, Wenxuan Guo, Tal Golan, Nikolaus Kriegeskorte

**Affiliations:** Department of Psychology, University of Cambridge, Cambridge, UK; Cognition and Brain Sciences Unit, University of Cambridge, Cambridge, UK; Department of Experimental Psychology, Justus Liebig University, Giessen, Germany; Centre for Mind, Brain and Behaviour (CMBB), University of Marburg and Justus Liebig University Giessen, Germany; Zuckerman Mind Brain Behavior Institute, Columbia University, New York, USA; Department of Psychology, Columbia University, New York, USA; Department of Neuroscience, Columbia University, New York, USA; Department of Electrical Engineering, Columbia University, New York, USA

**Keywords:** face perception, face similarity, face identification, Basel Face Model, deep neural networks

## Abstract

Human vision is attuned to the subtle differences between individual faces. Yet we lack a quantitative way of predicting how similar two face images look, or whether they appear to show the same person. Principal-components-based 3D morphable models are widely used to generate stimuli in face perception research. These models capture the distribution of real human faces in terms of dimensions of physical shape and texture. How well does a “face space” defined to model the distribution of faces as an isotropic Gaussian explain human face perception? We designed a behavioural task to collect dissimilarity and same/different identity judgements for 232 pairs of realistic faces. The stimuli densely sampled geometric relationships in a face space derived from principal components of 3D shape and texture (Basel Face Model, BFM). We then compared a wide range of models in their ability to predict the data, including the BFM from which faces were generated, a 2D morphable model derived from face photographs, and image-computable models of visual perception. Euclidean distance in the BFM explained both similarity and identity judgements surprisingly well. In a comparison against 14 alternative models, we found that BFM distance was competitive with representational distances in state-of-the-art image-computable deep neural networks (DNNs), including a novel DNN trained on BFM identities. Models describing the distribution of facial features across individuals are not only useful tools for stimulus generation. They also capture important information about how faces are perceived, suggesting that human face representations are tuned to the statistical distribution of faces.

## Introduction

Recognizing people by their faces is crucial to human social behaviour. Despite much work on the neural and behavioural signatures of face perception (see e.g. (1–4)), there is currently no quantitative model to predict how alike two faces will look to human observers. Advances in deep learning have yielded powerful artificial systems for face and object recognition (5–7), and 3D modelling and rendering techniques make it possible to systematically explore the space of possible faces (8–10). Here we comprehensively sample large sets of realistic face pairs from one such generative face model. We investigate how well its statistically-derived face space predicts perceived dissimilarity among faces, compared to a wide range of alternative models.

Since faces of different people are structurally highly similar and vary along continuous dimensions (nose length, jaw width, etc.), it is helpful to think of faces as forming a continuous “face space” (11–13). A face space is an abstract space in which each face occupies a unique position, and the dimensions span the ways in which physiognomic features can vary between faces. The origin of the multidimensional space is often defined as the average face: the central tendency of the population of all faces, or, for an individual, the sample of faces encountered so far. There is perceptual and neural evidence that face encoding adapts to better distinguish among the faces we encounter more frequently either over the lifetime or within the recent experience (2, 14–16). In a computational model, there are many conceivable ways one could parameterise and span the space of possible faces. For example, deep neural networks (DNNs) trained on face recognition represent individual faces in terms of combinations of complex non-linear image features (e.g. (17–19)). On the other hand, 3D morphable models represent faces in terms of a geometric mesh defining the face’s shape, and a texture map defining the colouration at each point on the face (8, 9, 20). 3D morphable models are very useful tools in face perception research because they can be used to generate novel realistic faces for which ground-truth properties are known, and which can be rendered under carefully-controlled viewing conditions (8–10, 15, 21).

We used the Basel Face Model (BFM) (8), a widely used 3D morphable model in both computer graphics and face perception research (e.g. (22, 23)). The BFM is a 3D generative graphics model that produces nearly photorealistic face images from latent vectors describing shape and texture of the surfaces of natural faces. The model is based on principal component analysis (PCA) of 3D scans (acquired using coded light system) of 200 adult faces (8). We used the BFM both as a stimuli generator and as a candidate model of facial dissimilarity. One appealing aspect of 3D morphable models like the BFM is that they provide a “ground truth” about the similarity relationships between different faces, allowing researchers to systematically vary similarity. However, distances within the BFM are defined in units of standard deviation within the sample of scanned faces. Units are therefore a statistical measure of how much facial features vary across real individuals. It is therefore not intuitively obvious that distances within a PCA-based space should predict perceived similarity well.

The goal of our study was to better understand perceived face similarity by comparing it to diverse candidate representational spaces. We are particularly interested in how perceived similarity relates to the statistics of how faces vary across individuals. We therefore chose to systematically sample different face relationships in a “face space” in which one unit of distance along each dimension captures one standard deviation of face structure/texture variation in a large sample of individuals. Is perception sensitive to the statistical distribution of face variation captured in this model, or is perceived similarity dominated by simpler considerations like how similar two faces are as 3D geometries or 2D images? The idea that face perception is “statistically tuned” has long been suggested, in the neurophysiological (15) and psychological literature (11), so this is an important question to ask. Our paper reports the first comprehensive test of a statistical face-space model of human face similarity percepts, by measuring how well distances in face space predict a rich dataset of human similarity and identity judgements. We compare the statistical face-space model to a large set of alternative metrics and models, which enables us to evaluate a wider range of hypotheses. Only four out of the 15 models we compared (pixel, VGG-Object, VGG-Face, and Active Appearance Model) were considered in previous works (10, 21, 24). Apart from low-level image descriptors and deep neural network models, we also included a 2D morphable model (25, 26) and models that capture “mid-level” information, for example about the 3D shape of a face. We perform the most comprehensive model comparison to date, comparing a total of 15 diverse models spanning DNNs, lower-level models, Eigenfaces, the Active Appearance Model, and models based on BFM geometry and facial coordinates.

To gain more insight into how humans judge face dissimilarity and identity, we designed a novel task for efficiently obtaining high-fidelity dissimilarity and identity judgements. Previous studies used multi-arrangement task for individual images of objects (27). Here, we introduce a multi-arrangement task for pairs of images. We selected stimulus pairs to systematically sample geometric relations in statistical face space, exhaustively creating all combinations from a large set of facial vector lengths and angles (Figure 1b and 2a), see Methods). This experimental design allowed us to carefully test how well the BFM’s statistical space captured human judgements, and what the geometric relationship was between BFM distances and distances in human perceptual space. We are not seeking the best applied model for human perception of real-world photographic faces. Instead, we want to better understand the perceptual metric of “face similarity”, by comparing it to diverse quantitative measures of face similarity, enabled by the usage of generated faces. Therefore, we trade photo-realism and real-world variability for a densely, systematically sampled face space where distances correspond to statistical variations in facial shape and texture. Having a controlled stimulus set enabled us to test more models and model types that otherwise would not be possible to consider such as shape, texture, and geometric relationships in the 3D morphable model. This sampling procedure also made it possible to probe isotropy. A representational space is perceptually isotropic if perceived dissimilarity remains constant as the direction of the pair of face vectors is rotated in any direction around the origin (while preserving their lengths and angle).

**Fig. 1.**
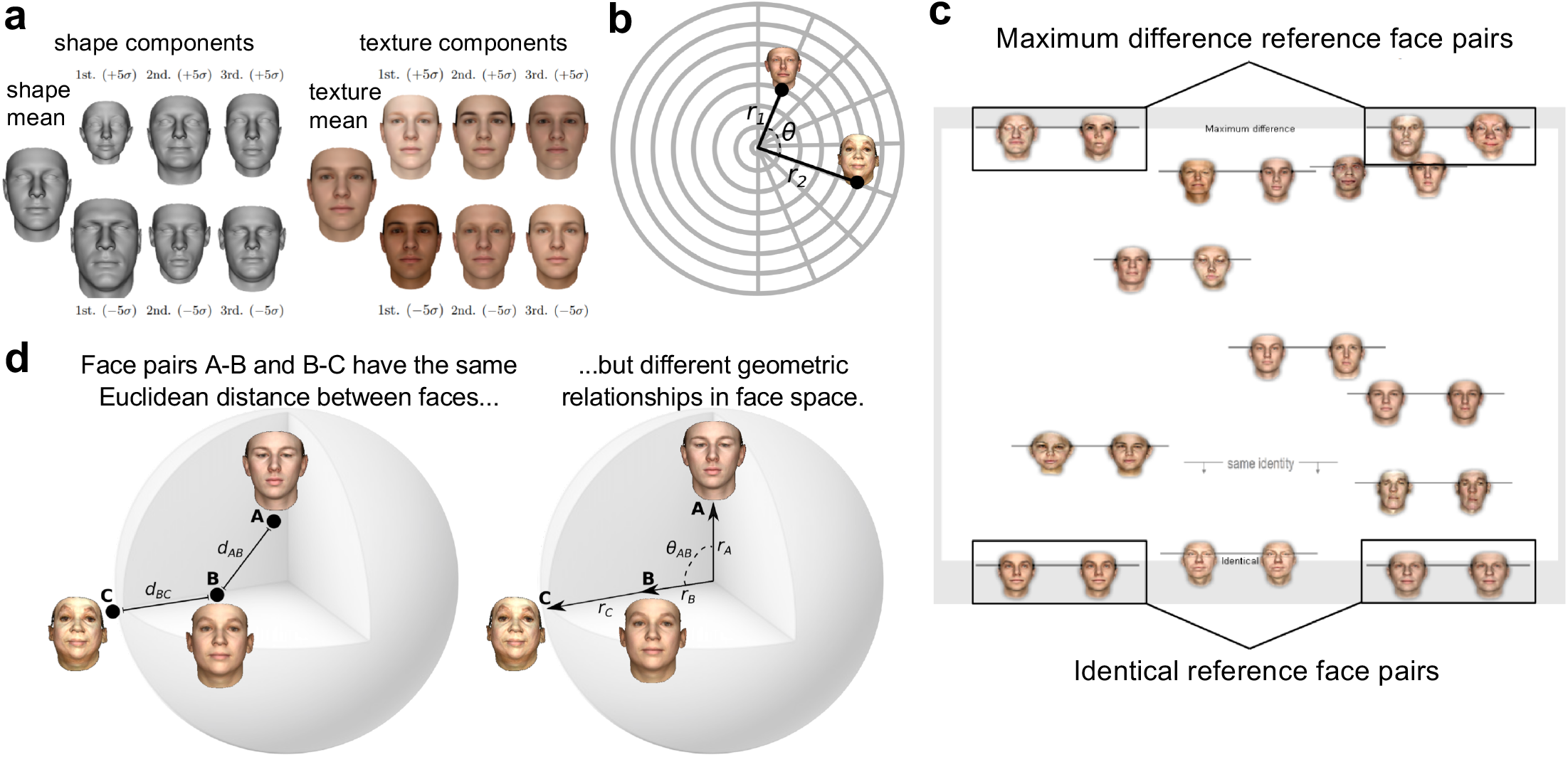
Selecting pairs of faces from the BFM, and measuring perceived face dissimilarity and identity. **a)** Illustration of the generative Basel Face Model (BFM), in which faces are described by separate components specifying their 3D shape (left) and texture (right). Both shape and texture components have a mean shape or texture (leftmost column), that can be changed by manipulating principal components. The first three PCs within each subspace are shown here (right three columns). The values of *σ* (-/+ 5*σ*) mean going a number of standard deviations away from the mean face in the direction of a given PC. Reproduced with permission from (8). **b)** Stimulus selection. We defined stimuli as pairs of vectors in the BFM with radial lengths *r*_1_ and *r*_2_, and angle between them (*θ*). We sampled all unique combinations from eight *θ* values and eight radius values to obtain 232 face pairs. See Methods for details. **c)** Behavioural experiment task. Participants positioned the face pairs along the vertical axis of the screen according to their relative dissimilarity. Faces were arranged in random subsets of eight pairs, as in the example shown. The vertical position of the line linking face pairs determined their precise location. As points of reference for participants, two example face pairs depicting “Maximum difference” (top) and two example face pairs depicting “Identical” (bottom) were shown. For each subset, participants also placed a “same identity” bar indicating the point below which pairs of images appear to depict the same identity. **d)** Relationships between pairs of faces in a face space like the BFM can be thought of in terms of the Euclidean distance between them (left), or their geometric relationships relative to the centre of the face space (right). If perceived dissimilarity can be predicted from the Euclidean distance alone, then face pair AB should look exactly as similar to one another as face pair BC. However, if observers take the angular and radial geometry of face space into account, they may have substantially different similarities.

**Fig. 2.**
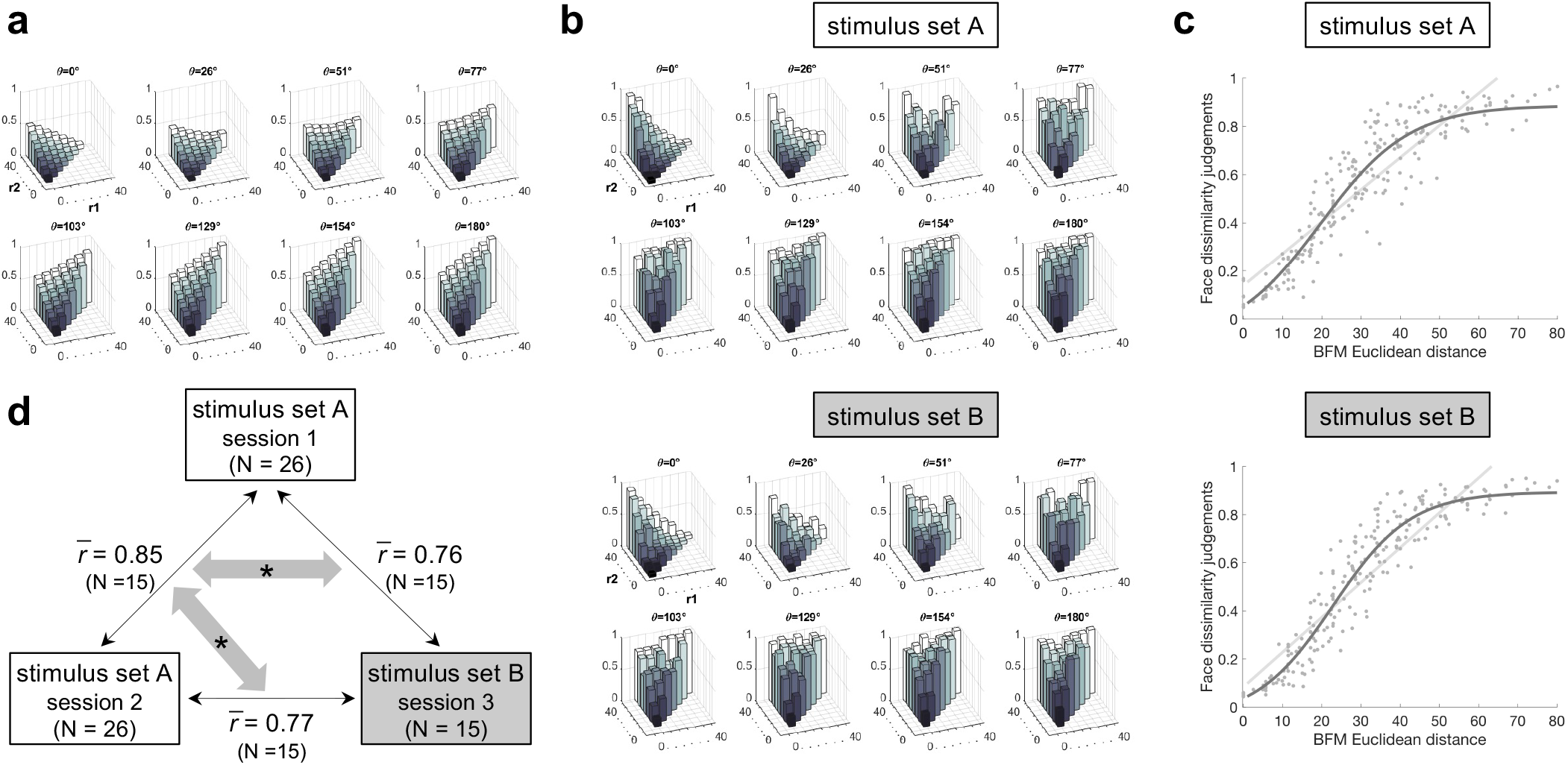
Face dissimilarity judgements as a function of distances in the BFM. **a)** Euclidean distances within the BFM for each face pair in the stimulus sets. Sets A and B had identical BFM geometries, but used different specific face exemplars. Each plot shows pairs of face vectors separated by a specific angle (*θ*), and arranged by the lengths of each of the two vectors (*radius*_1_ and *radius*_2_). Radii were sampled in eight evenly-spaced steps, from a length of zero (black), to a length of 40 units in the BFM (white). Normalised Euclidean distance between faces in each pair is indicated by the height of each bar, from zero (identical) to one (maximum distance within the stimulus set). **b)** Human face dissimilarity judgements for each face pair, for the stimulus set A (top) and B (bottom). Plotting conventions are as in 2a, except that bar height indicates human-rated dissimilarity, from zero (identical faces) to one (maximally dissimilar faces), averaged over participants and trials. **c)** Face dissimilarity judgements (y-axis) as a function of Euclidean distance in the BFM (x-axis) for the stimulus set A (top) and B (bottom). Each dot represents the mean dissimilarity rating for one pair of faces, averaged across participants and trials. The pale grey line represents the fit of a linear function and the dark grey line represents the fit of a sigmoidal function to the data. **d)** Replicability of face dissimilarity judgements between sessions 1 and 2 (using stimulus set A and the same participants), and session 3 (using stimulus set B and a subset of the participant group). All 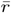 values are averages of between-session Pearson correlation coefficients across the subset of 15 participants that participated in all three sessions. Grey arrows with asterisks indicate significantly different correlations (one-tailed paired *t*-test across the 15 participants: *p* = 0.00000004, one-tailed paired Wilcoxon signed-rank test: *p* = 0.00003). The correlation of face dissimilarity judgements between the two sessions using stimulus set A (session 1 and 2) is higher than the correlation of judgments between stimulus sets (session 1 and session 3 or session 2 and session 3) for each of the 15 participants.

During the task, participants arranged pairs of face images on a large touch-screen according to how similar they appeared, relative to anchoring face pairs at the top (maximally different, diametrically opposed) and bottom (identical) of the screen and relative to other adjusted pairs (Figure 1c). This task has two advantages over standard pairwise dissimilarity ratings: (1) It provides a more fine-grained continuous measure of face dissimilarity within each pair (the vertical position at which the pair was placed on the screen). (2) The judgements are anchored not just to the extreme anchor pairs provided above and below the sorting arena, but also to the other adjustable pairs within each trial. Previous studies focused either on facial similarity or facial identity; by measuring both of them in the same task we are able to evaluate categorical same/different identity judgements in the context of continuous similarity judgements. We also tested whether the relative geometry within BFM was perceptually isotropic. Therefore, the stimulus set A and stimulus set B experiments had the same relative geometries but different face exemplars 1d. We sought to model both the continuous aspects of human face perception (graded dissimilarity) and its categorical aspects (same/different identity).

## Results

Participants (N=26) were highly reliable in their dissimilarity judgements using the novel arrangement task (mean correlation between participants = 0.80, mean correlation for the same participant between sessions = 0.85, the stimulus set A experiment). This provided a high-quality dataset with which to adjudicate between candidate models. We repeated the same experiment with a subset of the same participants (N=15) six months later, with a new independently sampled face set fulfilling the same geometric relations as the original stimulus set (stimulus set B experiment, see Methods). Participants in the stimulus set B experiment were also highly reliable in their dissimilarity judgements (mean correlation between participants = 0.79). This level of replicability allowed us to evaluate to what extent dissimilarity judgements depend on idiosyncrasies of individual faces, and to what extent they can be predicted from geometric relations within a statistical face space.

### Face dissimilarity judgements can be well predicted by distance in a statistical face space

We first asked how well human face dissimilarity judgements could be predicted by distances within the Basel Face Model (BFM), the principal-components-based face space from which our stimuli had been generated. Since we had selected face pairs to exhaustively sample different geometric relationships within the BFM, defined in terms of the angle between faces and the radial distance of each face from the origin, we were able to visualise human dissimilarity ratings in terms of these geometric features (Figure 2b). The patterns of human dissimilarity ratings closely resembled the patterns of Euclidean distances among our stimuli in the BFM space (Figure 2a). Given this, we plotted dissimilarity judgements for each face pair as a function of the Euclidean distance in the BFM (Figure 2c). To quantify how well the BFM approximates face dissimilarity judgements, we tested which functions best capture the relationship between behavioural dissimilarity judgements and BFM distances. We plotted the predictions of each fitted function over the data and compared their goodness of fit. If the BFM is a perfect approximator of face dissimilarity judgements a linear function would best describe the relationship between face dissimilarity judgements and the Euclidean distances in the BFM. We do not find this assumption to be completely true as the sigmoidal function better describes the relationship between face dissimilarity judgements and the Euclidean distances in the BFM (Figure 2c, the goodness of fit of linear function = 0.82, the goodness of fit of sigmoidal function = 0.86, one-sided Wilcoxon signed-rank test, *p <* 0.05). The sigmoidal relationship between the BFM and perceived distances suggests that observers have maximal sensitivity to differences between faces occupying moderately distant points in the statistical face space, at the expense of failing to differentiate between different levels of dissimilarity among very nearby or very far apart faces. This latter result may be related to the fact that faces with very large Euclidean distances in the BFM look slightly caricatured to humans (Supplementary Figure 1). We observed similar results in the stimulus set B experiment (face dissimilarity judgements using different face pairs with the same geometrical properties as in the stimulus set A experiment, see Methods for details, Figure 2c). This result suggests that the sigmoidal relationship between the BFM and perceived distances is observed regardless of the face pairs sampled. Overall, the BFM is a good, but not perfect, approximator of face dissimilarity judgements.

### Face identity judgements can be well predicted from the Euclidean distance in BFM

We also asked humans to judge whether each pair of faces depicted the same or different identity, and examined human identity thresholds in relation to the Euclidean distance between faces in the BFM. We found that moderately dissimilar faces in terms of the BFM Euclidean distance are often still perceived as having the same identity (Figure 3a). We observed similar results in the stimulus set B experiment (Figure 3a). This result may be related to humans having a high tolerance to changes in a personal appearance due to age, weight fluctuations, or skin complexion depending on the season.

**Fig. 3.**
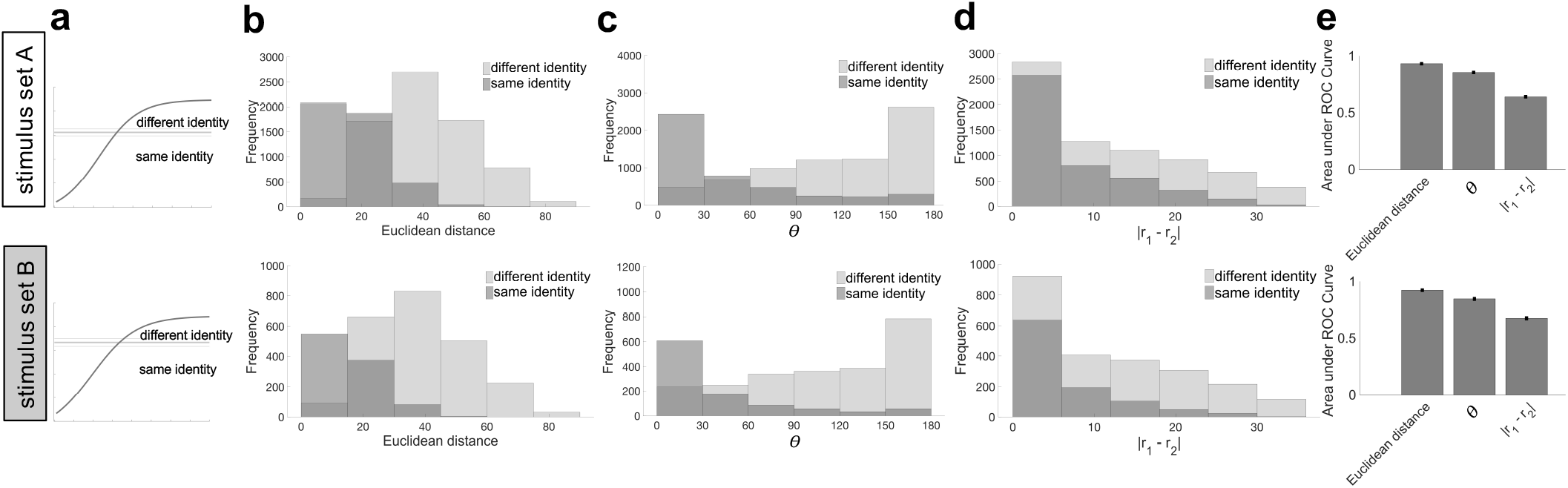
Identity judgements as a function of geometry within the BFM. **a)** Threshold for judging faces as belonging to the same/different identity, visualised relative to similarity judgements. The curved line shows the sigmoidal fit to face dissimilarity judgements (from Figure 2c) in the stimulus set A (top) and B (bottom). The thick horizontal line shows the mean placement of the “different identity” threshold bar, across participants and trials; thinner lines above and below indicate the standard error of the mean over participants. **b)** Histogram of how frequently face-pairs were judged as having the same identity (dark grey) or different identity (light grey), as a function of their Euclidean distance in the BFM. **c)** Histogram of same and different identity judgements as a function of angle (*θ*) between faces in the BFM. **d)** Histogram of same and different identity judgements as a function of the absolute difference between vector lengths in BFM (*r*_1_ and *r*_2_). **e)** Summary of how well each of the three BFM metrics in b-d discriminates face pairs judged as having the same vs different identity. Bars show the area under the receiver operating characteristic (ROC) curve calculated based on identity judgements using Euclidean distance, *θ*, and the absolute difference between *r*_1_ and *r*_2_.

Face pairs in the BFM can be analyzed in terms of their geometric characteristics relative to the centre of the face space or as the Euclidean distance between them. Therefore, we tested alternative predictors of face identity judgements: geometry in the BFM (*θ*, absolute difference between *r*_1_ and *r*_2_) and the Euclidean distance. We could predict whether two faces will be classified as the same individual by each of the predictors (Figure 3b, c, d). The Euclidean distance in the BFM predicted identity judgements marginally better than the angular and radial geometry of face space (Figure 3e).

### Relative geometry within BFM is approximately but not exactly perceptually isotropic

A representational space is perceptually isotropic if perceived dissimilarity remains constant as the direction of the pair of face vectors is rotated in any direction around the origin (while preserving their lengths and angle). We cannot test for isotropy in the stimulus set A experiment, because for each geometric relationship ((*θ*), r1, r2) we only have one sample of face pairs. To address this limitation, we compared the responses to the stimulus set A with either a repeated measurement of the responses to the same stimulus set or with the responses to stimulus set B, which had the same relative geometries but different face exemplars (i.e., each face pair had a different direction in BFM space). Participants completed two sessions of the stimulus set A experiment and a subset of participants (15 out of 26) completed the third session of the stimulus set B experiment. If the relative geometry within BFM is isotropic then the correlation between stimulus set A and B experiments should be the same as the correlation between two sessions of the stimulus set A experiment. The correlation between two sessions of the stimulus set A experiment for the subset of 15 participants is 0.85, the correlation between stimulus set A session 1 and stimulus set B session 3 experiment is 0.76, and the correlation between stimulus set A session 2 and stimulus set B session 3 experiment is 0.77 (Figure 2d). Replicability of face dissimilarity judgements between sessions 1 and 2 (using stimulus set A and the subset of 15 participants), and session 3 (using stimulus set B and a subset of the participant group) is significantly higher when the same stimulus set was used (one-tailed paired *t*-test across the 15 participants: *p* = 0.00000004, one-tailed paired Wilcoxon signed-rank test: *p* = 0.00003). The correlation of face dissimilarity judgements between the two sessions using stimulus set A (session 1 and 2) is higher than the correlation of judgments between stimulus sets (session 1 and session 3 or session 2 and session 3) for each of the 15 participants. These results suggest that the relative geometry within BFM is approximately, but not exactly, perceptually isotropic. However, since the stimulus set B experiment was performed six months after the stimulus set A experiment, the decreased correlation with stimulus set B could be attributed to the longer time between sessions with different face exemplars. Hence, we do not interpret the correlation difference as strong evidence against isotropy.

A related concept to isotropy is face space uniformity. The space is perceptually uniform if perceived dissimilarity remains constant as the pair of face points is translated anywhere in the space. This is a looser requirement than isotropy because all it preserves is the Euclidean distance between face points, not their geometry relative to the origin. For example, if the space is uniform, then it should not matter whether one face is at the origin and the other is 10 units away, or both faces are 5 units away from the origin in opposite directions. We searched for evidence of perceptual uniformity in the stimulus set A experiment, by binning face pairs into groups with similar Euclidean distance and then evaluating whether the angle between the faces explains variance in perceived dissimilarity. If the space is non-uniform, we might expect faces with larger angular differences to appear more different, even if they have identical Euclidean distance. We find only weak evidence for any non-uniformity in the face space (Supplementary Figure 2).

### Deep neural networks including a novel BFM-identity trained DNN predict well perceived face dissimilarity

After establishing that face dissimilarity judgements can be predicted from the Euclidean distance relatively well, we wanted to test a wide range of models to examine whether there is one model that best explains face dissimilarity judgements or there are multiple models that can explain the data equally well.

All models tested are schematically presented in Figure 4a. We considered Euclidean or Cosine distances within the full BFM space. Simple alternative models consisted of a 3D mesh model, RGB pixels, GIST, Eigenfaces, and face configurations (“0th order” configuration (location of 30 key points such as eyes, nose, mouth), “1st order” configuration (distances between key points), and “2nd order” configuration (ratios of distances between key points)). We also considered a 2D morphable model, namely the Active Appearance Model (25), trained to summarise facial shape and texture of natural faces (26). Finally, the last class of models consisted of deep neural networks (DNNs) of either a 16-layer VGG (28) or 8-layer Alexnet architecture (29). VGG was trained either to recognise objects, identify faces from real-world photographs, or to recognise synthetic identities generated from the BFM. All DNNs were trained on recognition tasks rather than to report dissimilarity directly. Predicted dissimilarities of each face pair for each model are shown in Supplementary Figure 3).

**Fig. 4.**
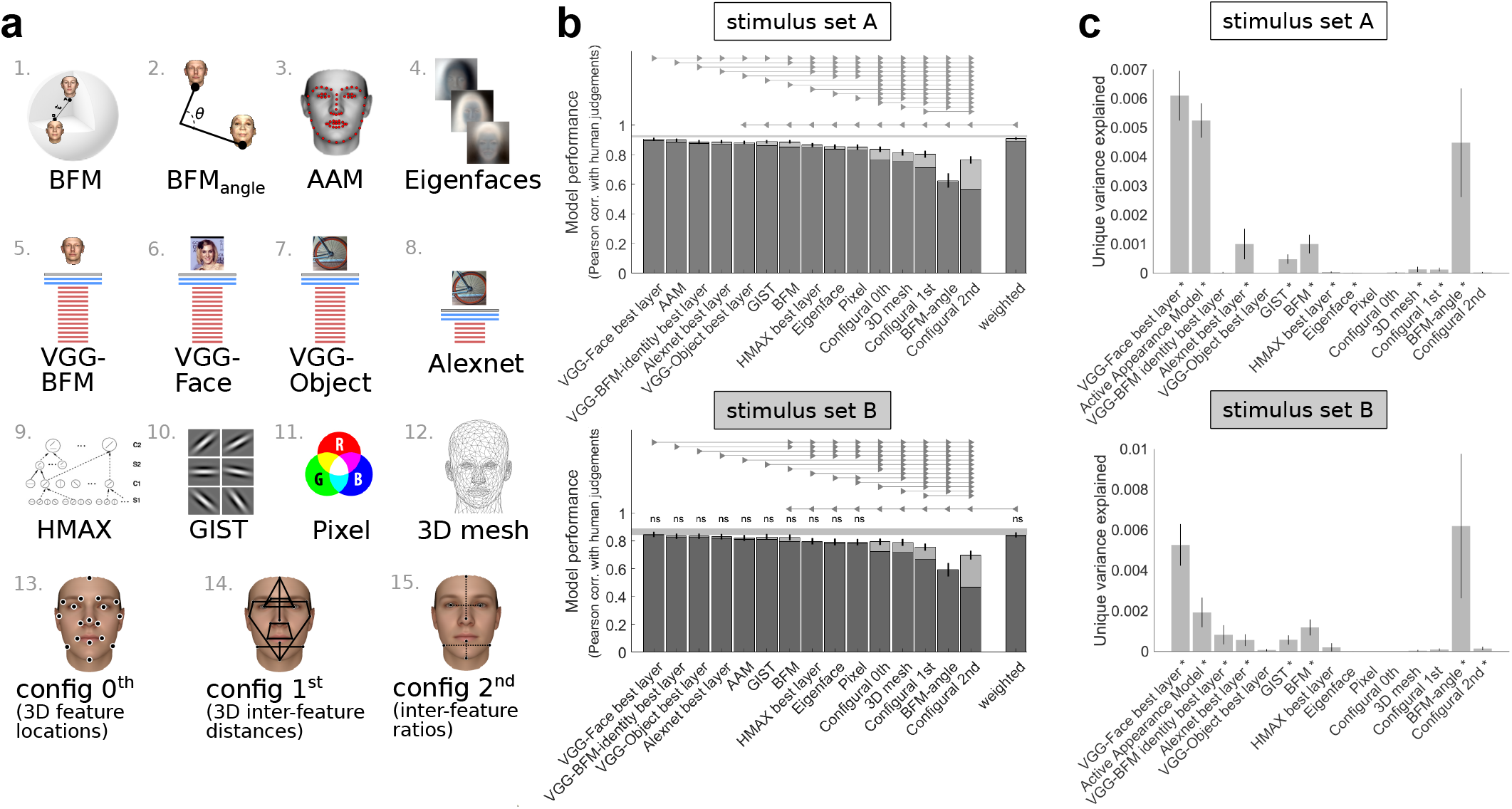
Comparing diverse models in their ability to predict face dissimilarity judgements. **a)** Schematic illustration of models compared (see 1 for full details). Two models were based on the 3D morphable model from which faces were generated: 1. Euclidean distance in the 3D morphable model and 2. Cosine distance within the full BFM coordinate space. Other models included 3. Active Appearance Model (AAM) and 4. Eigenfaces. Deep neural network (DNN) models consisted of a 16 layer VGG architecture trained on either (5.) BFM faces, (6.) face photographs or (7.) objects, and (8.) an 8-layer Alexnet architecture trained on objects. Alternative models were: 9. a shallower HMAX neural network; 10. GIST image descriptors; 11. raw pixel values; 12. raw 3D face mesh; and configural models: 13. “0th order” configuration (location of 30 key points such as eyes, nose, mouth); 14. “1st order” configuration (distances between key points); and “2nd order” configuration (ratios of distances between key points). **b)** Ability of each model to predict face dissimilarity judgements in the stimulus set A (top) and B (bottom). Bars show a Pearson correlation between human-judged face dissimilarity and face-pair distance within each model. The dark lower region of each bar shows performance for raw model distances, while the paler upper region shows additional performance gained if model distances are transformed by a compressive nonlinearity (a sigmoidal function fitted to data from training participants and face-pair stimuli). All models were significantly correlated with human data (*p <* 0.05 corrected). The grey bar represents the noise ceiling, which indicates the expected performance of the true model given the noise in the data. The final bar shows the performance of a linear weighted combination of all models, fitted using non-negative least-squares. Fitting of sigmoidal transforms and linear reweighting was performed within the same cross-validation procedure, fitting and evaluating on separate pools of both participants and stimuli. Error bars show the standard error of the mean (95% confidence interval over 1,000 bootstrap samples). Horizontal lines show pairwise differences between model performance (*p <* 0.05, Bonferroni corrected across all comparisons). Models connected by triangular arrow markers indicate a significant difference, following the convention in (38), with the direction of the arrow marker showing which model is superior. All statistical tests shown were performed on the raw untransformed version of each model. **c)** Unique variance in face dissimilarity judgements computed using a hierarchical general linear model (GLM) for the stimulus set A (top) and B (bottom). For each model, unique variance is computed by subtracting the total variance explained by the reduced GLM (excluding the model of interest) from the total variance explained by the full GLM, using non-negative least squares to find optimal weights. Models that explain significant unique variance are indicated by an asterisk (one-sided Wilcoxon signed-rank test, *p <* 0.05 corrected). Error bars show the standard error of the mean based on single-participant unique variance.

We inferentially compared each model’s ability to predict face dissimilarity judgements, in both their raw state and after fitting a sigmoidal transform to model-predicted dissimilarities, using a procedure cross-validated over both participants and stimuli (see Methods). The highest-performing model was the VGG deep neural network trained on face identification (VGG-Face (17)); Figure 4b). We included in the comparison the highest performing layer of each deep neural network (for the comparison of all VGG layers see Supplementary Figure 4). To remove the mismatch between training (real naturalistic faces) and test distributions of faces (BFM), we trained the same VGG-16 architecture to classify identities sampled from the Basel Face Model 2019 (see Methods). The best layer of a VGG trained on BFM identities does not perform better than the best layer of a VGG trained on natural faces, and does not capture any unique variance 4c). In all three VGG training scenarios (objects, natural faces, and BFM faces), match to human judgements peaked in late convolutional layers, before declining again in fully-connected layers (for the comparison of all VGG-BFM-identity layers see Supplementary Figure 5). Human face perception is captured by a DNN that has experience with real naturalistic faces; and there is no evidence that the performance of a DNN trained of real faces is diminished by testing it on generated faces. Several other models had high performances, and that of the 2D morphable Active Appearance Model (AAM) was not statistically different from that of VGG-Face. Euclidean distance within BFM was outperformed by VGG-Face and AAM, but not by any of the other models (Figure 4b, top). Performing the same analysis on the independent stimulus set B experiment revealed good reproducibility of the model rankings, even though the stimulus faces are different (Figure 4b, bottom). VGG-Face again achieved the highest performance, but in this dataset was not significantly superior to several other models: VGG-BFM-identity, VGG-Object, Alexnet, Active Appearance Model, and GIST. Again the BFM model was competitive with image-computable models, being outperformed only by VGG-Face and VGG-BFM-identity. Most models reached the noise ceiling in this second dataset, but this is likely because there was greater overall measurement noise, due to smaller sample size and one rather than two experimental sessions.

Having previously found that sigmoidally transforming BFM-predicted dissimilarities allowed them to better capture human perceived dissimilarities, we also evaluated all models after sigmoidally transforming their raw predictions (cross-validated, fitting and testing on separate participants and stimuli). Pale upper portions of bars in 4b indicate the performance of sigmoidally-fitted versions of each model, and Supplementary Figure 6 shows the result of statistical comparisons between them. All models better predicted human responses after fitting a sigmoidal function to their raw predicted distances, and produced a greater relative improvement for more poorly-performing models, but did not substantially affect model rankings (Figure 4b). VGG-Face predicts human judgements best, and BFM distance is competitive, being outperformed by no other models in the stimulus set B experiment, and only by VGG-Face in the stimulus set A experiment (see Supplementary Figure 6). Taking raw and sigmoidally-transformed performance across the two datasets into account, we found no single best model, but a set of consistently very highly performing ones. The four DNN models, the 3D morphable BFM, the 2D morphable AAM, and the image-statistic summary GIST model all excellently predicted face dissimilarity judgements. We observed that the average correlation between the models is 0.81 (with the highest correlation being 0.99 and the lowest correlation being 0.41, Supplementary Figure 9). These results mean that there is a different degree of similarity of model predictions between models tested.

In the supplementary analyses, we evaluate a number of other models based on the BFM: Euclidean and Cosine distances within the BFM shape dimensions only, BFM texture dimensions only, or within a four-dimensional subspace consisting of the dimensions capturing most variance in “person attributes” such as height, weight, age and gender (see Supplementary Figure 7). We also assessed models based on Euclidean distance within different numbers of principal components of the full BFM space (see Supplementary Figure 8). The performances are very similar for calculations performed on the full BFM space or on either the shape or texture subspaces using Euclidean distance or angle metrics (Supplementary Figure 7). The model with loadings on perceptually relevant dimensions of age, gender, weight, and height explained less variance than the full model (Supplementary Figure 7). Increasing the number of the principal component in the full model leads to a rapid increase in performance as the first 1-10 components are included (Supplementary Figure 8). The BFM model with 50 principal components (as used in (30)) is close to the performance of the full model. It seems that much smaller subspaces than the full 199-dimensional space may be sufficient to explain the variance in facial dissimilarity judgements.

There are substantial computational differences between the several models that all predict human perceived face dissimilarity well. Do they explain shared or unique variance in human judgements? To address this question we performed a unique variance analysis on all models. Several models explained a significant amount of unique variance, especially VGG-Face, BFM angle, and AAM models in both stimulus set A and stimulus set B experiments (Figure 4c). It is important to note, however, that the amount of unique variance explained by these models was very small.

If some models explain unique variance, perhaps combining them would explain more overall variance in face dissimilarity judgements? To address this question, we combined all models into one model via linear weighting, and asked whether this combined model explains more variance than each of the models alone. Model weights were assigned within the same procedure individual models were evaluated, cross-validating over both participants and stimuli. We found that in both datasets, the combined weighted model reached high performance, but did not exceed the performance of the best individual model (Figure 4b).

Models based on BFM or DNN feature spaces outperformed most others, including models based on the face perception literature (angle in the BFM “face space”, and simple configurations of facial features) and two baseline models (based on pixels or 3D face meshes). Poor performance of configural models is at odds with the previously proposed importance of inter-feature distances and ratios for face recognition (31–33). It seems that the richer combination of detailed facial landmarks along with their visual appearance as captured by the Active Appearance Model (26) is needed to well predict facial dissimilarity judgements. Such models are also well suited to describing variations across facial expressions (34). The success of the V1-like GIST model is surprising and may be due not to its unique explanatory power, but its high shared variance with more complex models for the image set used, although it is consistent with previous work finding that Gabor-based models explain variance in face matching experiments (35) and explain almost all variance in the face- and other complex shape-matching experiments when stimuli are tightly controlled (36). A person-attributes model, consisting only of the four dimensions which capture the highest variance (among the scanned individuals) in height, weight, age, and gender, did not perform well. This finding may seem surprising given that an earlier systematic attempt to predict face dissimilarity judgements from image-computable features found that dissimilarity was best predicted by weighted combinations of features that approximated natural high-level dimensions of personal characteristics such as age and weight (37). However, it seems that people use other or more than socially relevant dimensions when judging face dissimilarity in the experiment presented here.

## Discussion

We found that Euclidean distance in a principal-components-based 3D morphable model is a good approximator of human dissimilarity judgements. This is one of the few studies where such a model has been validated as providing quantitative predictions of perceived face dissimilarity and identity. The BFM was previously shown to capture face impressions (39) and personality traits (40) and different versions of face space models captured same-different face judgements (10), the similarity of randomly generated faces to four familiar identities (21), and facial dissimilarity judgements (24). The Basel 3D morphable model is derived from separate principal components analyses of 3D face structure and of facial texture and colouration. It is, therefore, a more sophisticated statistical model than earlier PCA-based face space models derived from 2D images, which only moderately well predicts face dissimilarity (41) and provides us with a theoretical understanding of how face perception can be mapped on principal component space. In our study, BFM has a dual role of being a good model and stimulus generator.

Face recognition requires us to solve all the usual problems of object recognition (lighting/view invariance), but it also has the more domain-specific challenge of representing the subtle structural differences between individual faces. It would make sense if our perception of these differences was attuned to the ways in which they vary in the natural population. In this paper, our emphasis is not on the broader challenges of object recognition (which any full model of facial similarity will also have to take into account, and which has received more research interest in the past), but specifically on measuring the perceptual importance of different face variations relative to a model of the true statistical variations among faces. This is why we opt to use very tightly controlled faces, in terms of pose and lighting, and comprehensively sample the face space. Exploring identity, view and lighting simultaneously is an important goal in the context of evaluating image-computable models of face perception. Using only frontal faces in a single lighting condition does make it harder to reject image-computable models, and this has to be taken into account in the interpretation. However, the BFM model, which is central to this study, by definition has perfect invariance to view and lighting, and so would have a trivial advantage over the image-computable models if view and lighting were varied. Our approach here gave image-based dissimilarity metrics their “strongest chance” – finding that distances in the BFM space better predicted human judgements, even under conditions favouring image metrics, demonstrates the importance of statistical feature deviation over and above mere visual difference between faces.

The success of the Euclidean distance alone to predict both dissimilarity and identity is striking, given that psychological face space accounts have assigned particular importance to the geometric relationships of faces relative to a meaningful origin of the space (2, 11, 12, 42, 43). For example, it has been reported that there are larger perceptual differences between faces that span the average face than not (44). Extensive behavioural and neuropsychological work has sought to relate the computational mechanisms underlying face perception to geometric relationships in neural or psychological face space. It has been proposed that face-selective neurons explicitly encode faces in terms of vectors of deviation from an average face, based on evidence from monkey electrophysiology (15, 30) and human psychophysical adaptation (2, 16, 45) although alternative interpretations of the latter have been made (46, 47). Our comprehensive sampling of face pairs with the full range of possible geometric relationships was tailored to reveal the precise manner in which distances from the origin, and angular distances between faces, affect perceived dissimilarity. Yet both dissimilarity and identity data were explained best by simply the Euclidean distance, with geometric relationships in face space accounting for no additional variance. Our results do not contradict previous studies, but suggest that effects of relative geometry may be more subtle than previously thought, when probed with large sets of faces that vary along diverse dimensions, rather than stimulus sets constructed to densely sample single or few dimensions (e.g. (43, 44)). Lastly, distances within BFM appear approximately but not exactly perceptually isotropic.

Distance within the BFM is not a perfect predictor of perceived dissimilarity. Firstly, like all morphable models, the BFM describes only the physical structure of faces, and so cannot account for familiarity, which we know to be substantial (cf. (42)). We did not explore people’s ability to parse structural differences between faces from sources of accidental differences between face images that are important for face recognition (lighting (48, 49) and viewpoint(48, 50)). Instead, we deliberately matched lighting and pose across faces in order to give image-based dissimilarity metrics their “strongest chance”. Finding that distances in the BFM space better predicted human judgements even, under conditions favouring image metrics, demonstrates the importance of statistical feature deviation over and above mere visual difference between faces. It is hard to predict how differences in lighting or viewpoint would affect the performance of the models but it may further differentiate wholly image-based models (e.g. GIST), from BFM that is perfectly view-invariant. DNNs lie on a continuum between image-dependent and invariant models as they learn partially view-invariant features, allowing them to robustly recognise faces under naturalistic viewing variations. A second reason that the BFM distance is imperfect as a dissimilarity predictor is that there remains unexplained variance that is reliable across individual observers but not captured by any version of the BFM model. There are several possible reasons for this. The BFM has limitations as a 3D morphable model, for example, it is based on the head scans of only 200 individuals, and this sample is biased in several ways, for example towards white, relatively young, faces. The sub-optimal performance could also be due to fundamental limitations shared by any physically-based model (42), such as its inability to capture perceptual inhomogeneities relating to psychologically relevant distinctions such as gender, ethnicity, or familiarity. It would be interesting to test in the future whether newer 3D morphable face space models capture more of the remaining variance in human dissimilarity judgements (9). The task presented here provides an efficient way to test the perceptual validity of future face space models.

We found that humans often classify pairs of images as depicting the same identity even with relatively large distances in the BFM. Two faces may be perceptibly different from one another, while nevertheless appearing to be “the same person”. The ability of the visual system to generalise identity across some degree of structural difference may be analogous to invariance to position, size and pose in object recognition (51). Face images generated from a single identity form a complex manifold as they may vary in age, weight, expression, makeup, facial hair, skin tone colour, and more (9). Given that we need to robustly recognise identity despite changes in these factors, it may not be surprising that there is a high tolerance for facial features when we judge one’s identity. The stimulus set contained very dissimilar faces, which provided an anchor for people’s definition of “different” and may influence moderately-dissimilar faces to look quite alike, in comparison. Participants seemed to interpret person identity quite generously, possibly imagining whether this face could be the same person if they aged, got tanned, or lost weight. “The same person” may be a not precisely defined concept, however people seem to agree what that concept means as they were consistent in judging the same/different identity boundary. Interestingly, the “different identity” boundary was close to the saturation point of face dissimilarity psychometric function. This result could be related to people dismissing all “different individuals” as completely different and focusing their fine gradations of dissimilarity only within the range of faces that could depict the same identity. The design of our current experiment required participants to locate a threshold of dissimilarity, above which they considered faces to be different identities. This means that identity and face dissimilarity judgements are entangled. It may be possible to dissociate them in future experiments by reporting identity independently from dissimilarity.

Our data show clearly that some models of face dissimilarity are worse than others. Simply taking the angle between faces in the BFM is a poor predictor, as is a set of higher-order ratios between facial features. Perhaps surprisingly, the model consisting only of the four dimensions which capture the highest variance in height, weight, age, and gender performed poorly. It is possible that this model may not have performed optimally because the face pair images in the face dissimilarity judgements might not have had enough variability across these dimensions. Age and gender were shown to explain variance in face MEG representations (52) and we show that they do explain variance in face dissimilarity judgements task, however to a lesser extent than better-performing models. It seems that people use other or more than socially relevant dimensions when judging face dissimilarity.

Among highly performing models, we found that several explain face dissimilarity judgements similarly well. One of the models that explains a surprisingly large amount of variance is GIST. It has been previously shown, that Gabor-based models explain face representations well (22, 35). The models compared contain quite different feature spaces. For example, object-trained and face-trained VGG models learn distinctly different feature detectors (6), yet explain a similar amount of variance in human face dissimilarity judgements. Both object-trained and face-trained VGG models also explain a similar amount of variance in human inferior temporal cortex (53), and object-trained VGG explains variance in early MEG responses (54). The “face space” within a face-trained DNN organises faces differently than they are arranged in the BFM’s principal components, for example, clustering low-quality images at the “origin” of the space, eliciting lower activity from all learned features (42).

It is perhaps remarkable that distances within the BFM are approximately as good at capturing perceived face dissimilarities as image-computable DNNs. Distances within the BFM contain no information about either the specific individuals concerned, or the image-level differences between the two rendered exemplars. DNNs, on the other hand, are image-computable and thus capture differences between the visible features in the specific rendered images seen by participants. The high success of the relatively impoverished BFM representation may highlight the importance of statistical face distributions to human face perception. After all, the BFM simply describes the statistical dissimilarity between two faces, expressed in units of standard deviations within the sample of 200 head-scanned individuals. The power of this statistical description is consistent with previous evidence for the adaptability of face representations, coming from face aftereffects (2, 15, 16, 45), the “own-race effect” (55, 56), and inversion and prototype effects (14).

One of the reasons for the equally high performance of disparate models is that, for our stimulus set, several models made highly correlated predictions (Figure 9), making it difficult to discriminate between them based on the current data. Model dissociation was also found to be difficult when studying the representations of face dissimilarity in the human fusiform face area, where the Gabor filters model performed similarly to the face space sigmoidal ramp-tuning model (22). Stimulus optimisation methods could be used in the future to identify sets of stimuli for which current well-performing candidate models make maximally dissimilar predictions (38, 57, 58).

Note that the models included here are distributed over a vast space and so we cannot in a single study vary each of their properties. The BFM and image-computable models have different advantages and disadvantages for predicting human dissimilarity judgements. The BFM model has privileged access to distances in a shape and texture space normalized to represent the distribution of natural faces. However, the image-computable models have privileged access to image features of the kind known to be represented throughout the ventral visual stream. Similarly, the 3D mesh model has privileged access to the shape of the face surfaces. Comparing many models provides constraints for computational theory, but it will take many studies to drive the field toward a definitive computational model of face perception.

We conclude that as well as being a useful tool for generating faces in face perception research, principal-components based generative models capture important information about how faces are represented in human perception.

## Methods

### Stimuli

Each face generated by the BFM corresponds to a unique point in the model’s 398-dimensional space (199 shape dimensions, and 199 texture dimensions), centred on the average face. The relative locations of any pair of faces can therefore be summarised by three values: the length of the vector from the origin to the first face *r*_1_, the length of the vector from the origin to the second face *r*_2_, and the angle between the two face vectors *θ* (see Figure 1b). To create a set of face pairs spanning a wide range of relative geometries in face space, we systematically sampled all pairs of eight possible radius values combined with eight possible angular values. Possible angular values were eight uniform steps between 0 and 180 degrees, and possible vector lengths were eight uniform steps between 0 and 80 units in the BFM. The measure of 80 units corresponds to 80 standard deviations. We chose image pairs that were far enough in the BFM but they did not look very caricatured (Supplementary Figure 1). For all eight angles and eight eccentricities, there were 232 unique face relationships when considering (a) that exchanging the two radii yields the same relationship for a given angle and (b) that the angle is irrelevant when one of the radii is 0. When both radii are >0, there are (7[non-zero radii] * 7[non-zero radii] + 7)/2 = 28 pairs of nonzero radius combinations (including identical radii), and so there are 28 * 8[angles] = 224 relationships between faces. When one of the radii is 0, then the angle is irrelevant, and there are an additional eight radius combinations (the other radius can take each of the eight values). For each relative geometry, we then sampled two random points in the full 398-dimensional BFM space that satisfied the given geometric constraints. We generated two separate sets of face pairs with the same relative geometries but different face exemplars, by sampling two independent sets of points satisfying the same geometric constraints. The two sets (stimulus set A and stimulus set B) were used as stimuli in separate experimental sessions (see “Psychophysical face pair-arrangement task”).

### Participants

Human behavioural testing took place over three sessions. Twenty-six participants (13 female) took part in sessions 1 and 2, and a subset of 15 (6 female) took part in session 3. All testing was approved by the MRC Cognition and Brain Sciences Ethics Committee and was conducted in accordance with the Declaration of Helsinki. Volunteers gave informed consent and were reimbursed for their time. All participants had a normal or corrected-to-normal vision.

### Psychophysical face pair-arrangement task

The procedure in all sessions was identical, the only difference being that the same set of face pair stimuli was used in sessions 1 and 2, while session 3 used a second sampled set with identical geometric properties. Comparing the consistency between sessions 1, 2, and 3 allowed us to gauge how strongly human judgements were determined by geometric relationships in face space, irrespective of the individual face exemplars.

During an experimental session, participants were seated at a comfortable distance in front of a large touch-screen computer display (43” Panasonic TH-43LFE8-IR, resolution 1920×1080 pixels). On each trial, the participant saw a large white “arena”, with a randomly arranged pile of eight face pairs in a grey region to the right-hand side (see Figure 1c). The two faces within each pair were joined together by a thin bar placed behind the faces, and each pair could be dragged around the touch-screen display by touching. Each face image was rendered in colour with a transparent background and a height of 144 pixels (approximately 7.1cm on screen).

The bottom edge of the white arena was labelled “Identical” and the top edge was labelled “Maximum difference”. Two example face pairs were placed to the left and to the right of the “Identical” and “Maximum difference” labels to give participants reference points on what identical and maximally different faces look like. The maximally different example faces had the largest geometric distance possible within the experimentally sampled geometric relationships (i.e. the Euclidean distance in the BFM = 80) in contrast to identical faces (i.e. the Euclidean distance in the BFM = 0). The same example pairs were used for all trials and participants.

Participants were instructed to arrange the eight face pairs on each trial vertically, according to the dissimilarity of the two faces within the pair. For example, two identical faces should be placed at the very bottom of the screen. Two faces that look as different as faces can look from one another should be placed at the very top of the screen. Participants were instructed that only the vertical positioning of faces would be taken into account (horizontal space was provided so that face pairs could be more easily viewed, and so that face pairs perceived as being equally similar could be placed at the same vertical location). On each trial, once the participant had finished vertically arranging face pairs by dissimilarity, they were asked to drag an “identity line” (see Figure 1c) on the screen to indicate the point below which they considered image pairs to depict “the same person”. Once eight face pairs and the identity line were placed, participants pressed the “Done” button to move to the next trial. Each session consisted of 29 trials.

### Representational similarity analysis

We used representational similarity analysis (RSA) to evaluate how well each of a set of candidate models predicted human facial (dis)similarity judgements (59). For every model, a model-predicted dissimilarity was obtained by computing the distance between the two faces in each stimulus pair, within the model’s feature space, using the model’s distance metric (see “Candidate models of face dissimilarity”). Model performance was defined as the Pearson correlation between human dissimilarity judgements and the dissimilarities predicted by the model. We evaluated the ability to predict human data both for each individual model and for a linearly weighted combination of all models. To provide an estimate of the upper bound of explainable variance in the dataset, we calculated how well human data could be predicted by data from other participants, providing a “noise ceiling”.

Noise ceilings, raw model performance, sigmoidally-transformed model performance, and reweighted combined model performance were all calculated within a single procedure, cross-validating over both participants and stimuli (60). On each of 20 cross-validation folds, 5 participants and 46 face pairs were randomly assigned as test data, and the remaining stimuli and participants were used as training data. On each fold, a sigmoidally-transformed version of each model was created, by fitting a logistic function to best predict dissimilarities for training stimuli, averaged over training participants, from raw model distances. Also on each fold, a reweighted combined model was created using non-negative least-squares to assign one positive weight to each of the individual models, to best predict the dissimilarity ratings for training stimuli, averaged over training participants. We then calculated, for each raw model, each sigmoidally transformed model, and for the combined reweighted model, the Pearson correlation with the model’s predictions for test stimuli for each individual test participant’s ratings. The average correlation over test participants constituted that model’s performance on this cross-validation fold. The upper bound of the noise ceiling was calculated within the same fold by correlating each test participant’s test-stimulus data with the average test-stimulus data of all test participants (including their own). The lower bound was calculated by correlating each test participant’s test-stimulus data with the average for all training participant’s test-stimulus data (60, 61). Means and confidence intervals were obtained by bootstrapping the entire cross-validation procedure 1,000 times over both participants and stimuli. We first determined whether each model was significantly different from the lower bound of the noise ceiling, by assessing whether the 95% confidence interval of the bootstrap distribution of differences between model and noise ceiling contained zero (60, 61), Bonferroni corrected for the number of models. Models that are not significantly different from the lower bound of the noise ceiling can be considered as explaining all explainable variance, given the noise and individual differences in the data. We subsequently tested for differences between the performance of different models. We defined a significant pairwise model comparison likewise as one in which the 95% confidence interval of the bootstrapped difference distribution did not contain zero, Bonferroni corrected for the number of pairwise comparisons.

### Area under the receiver operating characteristic curve calculation

We calculate the Receiver Operating Characteristic (ROC) curve such that for each Euclidean distance, (*θ*) or absolute difference between radii we first run a logistic regression (classifier) to predict identity (based on position relative to identity line). Second, we compute ROC curve on the classifier output. Specifically, for a range of discriminatory thresholds (0 to 1), we compute the true positive rate and false positive rate. ROC is the plot of true positive rate vs false positives rates. The Area Under the Curve (AUC) represents how well the classifier performs at various threshold settings. The threshold is the value at which the classifier will assign a label to a given input, by comparing the probability of that input vs the threshold.

### Unique variance analysis

We used a general linear model (GLM) to evaluate unique variance explained by the models (62). For each model, unique variance was computed by subtracting the total variance explained by the reduced GLM (excluding the model of interest) from the total variance explained by the full GLM. For model m, we fit GLM on X = “all models but m” and Y = data, then we subtract the resulting *R*^2^ from the total *R*^2^ (fit GLM on X = “all models” and Y = data). We performed this procedure for each participant and used non-negative least squares to find optimal weights. A constant term was included in the GLM model. We performed a one-sided Wilcoxon signed-rank test to evaluate the significance of unique variance contributed by each model across participants.

### Candidate models of face dissimilarity

We considered a total of 15 models of face dissimilarity in the main analyses (see Figure 4) and an additional 3 models in supplementary analyses (see Supplementary Figure 7). Each model consists of a set of features derived from either a face image, BFM coordinates, or 3D mesh, combined with a distance metric (see Table 1).

**Table 1.**
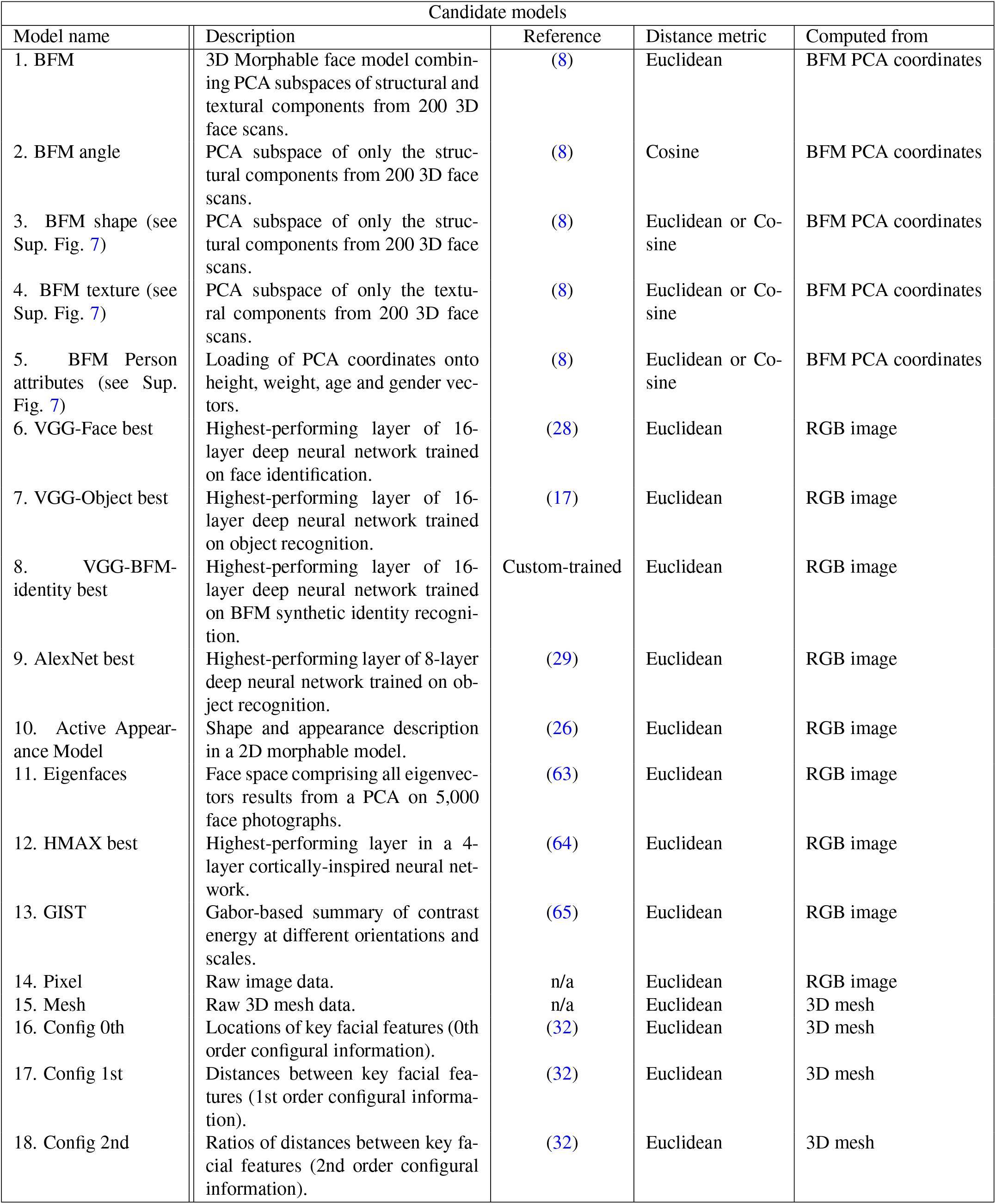
Candidate models of face dissimilarity.

#### Basel Face Model

We considered two variant models based on the principal-component space provided by the BFM: (1) “BFM Euclidean” took the Euclidean distances between faces in the full 398-dimensional BFM space and (2) “BFM angle” which took the cosine distance between face vectors in the full 398-dimensional space. For face pairs where cosine distance was undefined, because one face lay at the origin of BFM space, the angle between the two faces was defined as zero for the purposes of model evaluation.

To more fully explore the relationship between apparent dissimilarity and placements of faces in the full BFM space, we also considered linear and sigmoidal functions as candidates for predicting the relationship between the Euclidean distance in the BFM and face dissimilarity judgements. We estimated each model’s predictive performance as the Pearson correlation between the fitted model’s predicted dissimilarities and the dissimilarities recorded by the participant. We tested for significant differences between linear and sigmoidal function fits using a two-sided Wilcoxon signed-rank test. For each participant, we fitted the model to half of the data (session 1) and measured the predictive accuracy of the model in the second half of the data (session 2). The predictive accuracies were averaged across participants.

Finally, the BFM provides the axes onto which the height, weight, age, and gender of the 3D scanned participants most strongly loads. By projecting new face points onto these axes, we can approximately measure the height, weight, age and gender of each generated face. The “Person attributes” model took the Euclidean distance between faces, after projecting faces onto these four dimensions.

#### Active Appearance Model

Active Appearance Models (25) are 2D morphable models, generally trained on hand-annotated face images to identify a number of key facial landmarks from 2D photographs and describe visual structure around these landmarks. We applied a pretrained AAM provided in the Menpofit python package (https://www.menpo.org/menpofit/; (26)), which has been trained on 3,283 hand-labelled face photographs. This model describes 2D face structure in terms of the locations and appearance of 68 landmark features around the face border and internal features. After fitting the AAM to each stimulus face (100 fitting iterations) we appended the shape and appearance parameters to create a single vector describing the face, analogously to how shape and texture PCs were appended to create the full Basel Face Model descriptor. We then took the Euclidean distance between vectors as the predicted dissimilarity between faces in each stimulus pair.

#### Models based on 3D face structure

Face perception is widely thought to depend on spatial relationships among facial features (4, 31, 32, 66). We calculated the Euclidean distance between the 3D meshes that were used to render each face (“Mesh” model). We also used the geometric information within each face’s mesh description to calculate a first, second, and third-order configural model of facial feature arrangements, following suggestions by (32) and others (e.g. (31)) that face perception depends more strongly on distances or ratios of distances between facial features than raw feature locations. We selected 30 vertices on each face corresponding to key locations such as the centre and edges of each eye, the edges of the mouth, nose, jaw, chin, and hairline (see schematic in Figure 4a), using data provided in the BFM. The positions of these 30 vertices on each 3D face mesh formed the features for the “0th order” configural model. We then calculated 19 distances between horizontal and vertically aligned features (e.g. width of nose, length of nose, separation of eyes), which formed the “1st order” configural model. Finally, we calculated 19 ratios among these distances (e.g. the ratio of eye separation to eye height; the ratio of nose width to nose length), which formed the “2nd order” configural model.

#### Deep neural networks

We used a pre-trained state-of-the-art 16-layer convolutional neural network (VGG-16), trained on millions of images to recognize either object classes (28) or facial identities (17). Further details can be found in (17, 28). The dissimilarity predicted by DNN models was defined as the Euclidean distance between activation patterns elicited by each image in a face pair in a single layer. To input to DNN models, faces were rendered at the VGG network input size of 224×224 pixels, on a white background, and preprocessed to subtract the average pixel value of the network’s training image set.

We formed the VGG-BFM-identity classification network by training the VGG-16 architecture (28) (TorchVision’s implementation) to classify Basel Face Model 2019 face images of 8,631 identities. All of the images pertaining to one identity shared shape and texture latents (both randomly sampled from the Basel Face Model, once per identity), but had different expression latents, poses, lighting direction, and ambient lighting. We generated 363 images for each identity to roughly match the total number of training images in the VGGFace2 dataset (17). The rendered images were randomly cropped during training or centre cropped during validation, in both cases yielding an input image of 224 × 224 pixels. To further increase the images’ variability, we augmented the training examples using Albumentations (67). We only included naturalistic transformations such as grayscale transformation, brightness and contrast manipulations, noise addition, drop-out, grid distortion, and blurring. The input images were normalized by the channel-specific mean and standard deviation, computed from a subset of training images. We used PyTorch 1.7.1 and PyTorch Lightning 1.3.1 for model training. Models were trained for 30 epochs on minimizing the cross-entropy loss, using four GPUs. We used stochastic gradient descent with a weight decay of 0.0001, momentum of 0.9, and a minibatch size of 512. The initial learning rate was 0.01, and the step scheduler reduced the learning rate by a factor of 10 every 10 epochs. The model reached a 0.0 validation loss.

#### Low-level image-computable models

As control models, we also considered the dissimilarity of two faces within in terms of several low-level image descriptors: (1) Euclidean distance in raw RGB pixel space; (2) Euclidean distance within a “GIST” descriptor, image structure at four spatial scales and eight orientations (https://people.csail.mit.edu/torralba/code/spatialenvelope/); (3) Euclidean distance within the best-performing layer (C2) of HMAX, a simple four-layer neural network (http://cbcl.mit.edu/jmutch/hmin/), and (4) Euclidean distance within an “Eigenface” space consisting of the 4,999 dimensions obtained by running a Principal Component Analysis on the first 5,000 faces in the CelebA cropped and aligned dataset of celebrity faces (http://mmlab.ie.cuhk.edu.hk/projects/CelebA.html; (63)). To create the PCA space, photographs were resized to a height of 144 pixels and then cropped to the centre 144×144 pixels. For comparability with the images seen by participants, all low-level image-computable models operated on faces rendered on a white background at 144×144 pixel resolution.

## ACKNOWLEDGEMENTS

This research was supported by the Wellcome Trust [grant number 206521/Z/17/Z] awarded to KMJ; the Alexander von Humboldt Foundation postdoctoral fellowship awarded to KMJ; the Alexander von Humboldt Foundation postdoctoral fellowship awarded to KRS; the Wellcome Trust and the MRC Cognition and Brain Sciences Unit. For the purpose of open access, the author has applied a CC BY public copyright licence to any Author Accepted Manuscript version arising from this submission.

## AUTHOR CONTRIBUTIONS

JOK, KMJ and NK designed the experiments. KMJ collected the data. KMJ, JOK and KRS performed the analyses. WG and TG developed a VGG-BFM-identity model. KMJ and KRS wrote the paper. All authors edited the paper. NK supervised the work.

## DATA AND CODE AVAILABILITY

The datasets and code generated during the current study are available from the corresponding author on request.

## Supplementary Note 1: Appendix

**Supplementary Fig. 1.**
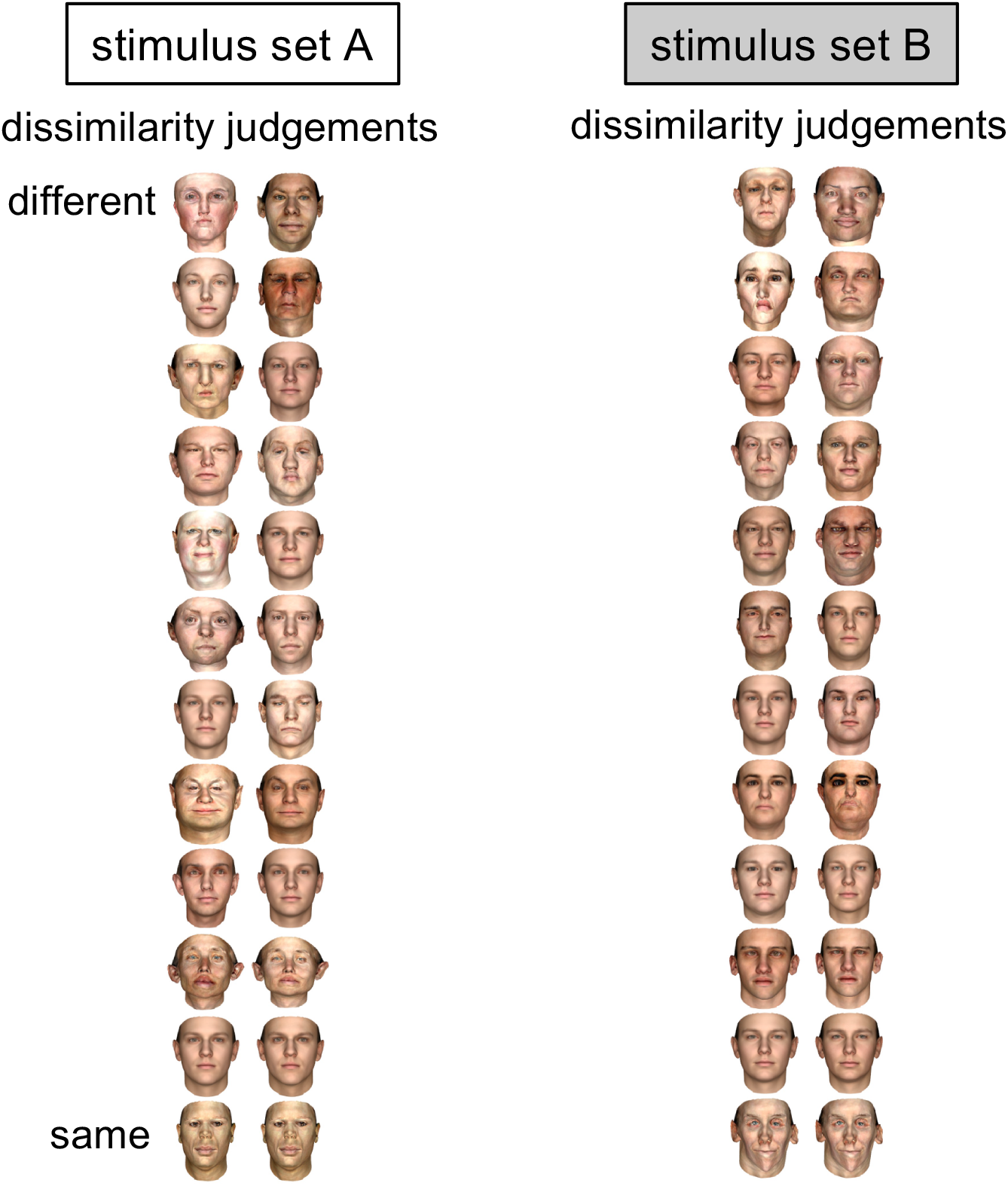
Face dissimilarity rankings by humans. Columns display face pairs according to their average rated dissimilarity by humans from most dissimilar (top) to most similar (bottom), visualising every 20th face pair in each ranked set within stimulus sets A (left) and B (right).

**Supplementary Fig. 2.**
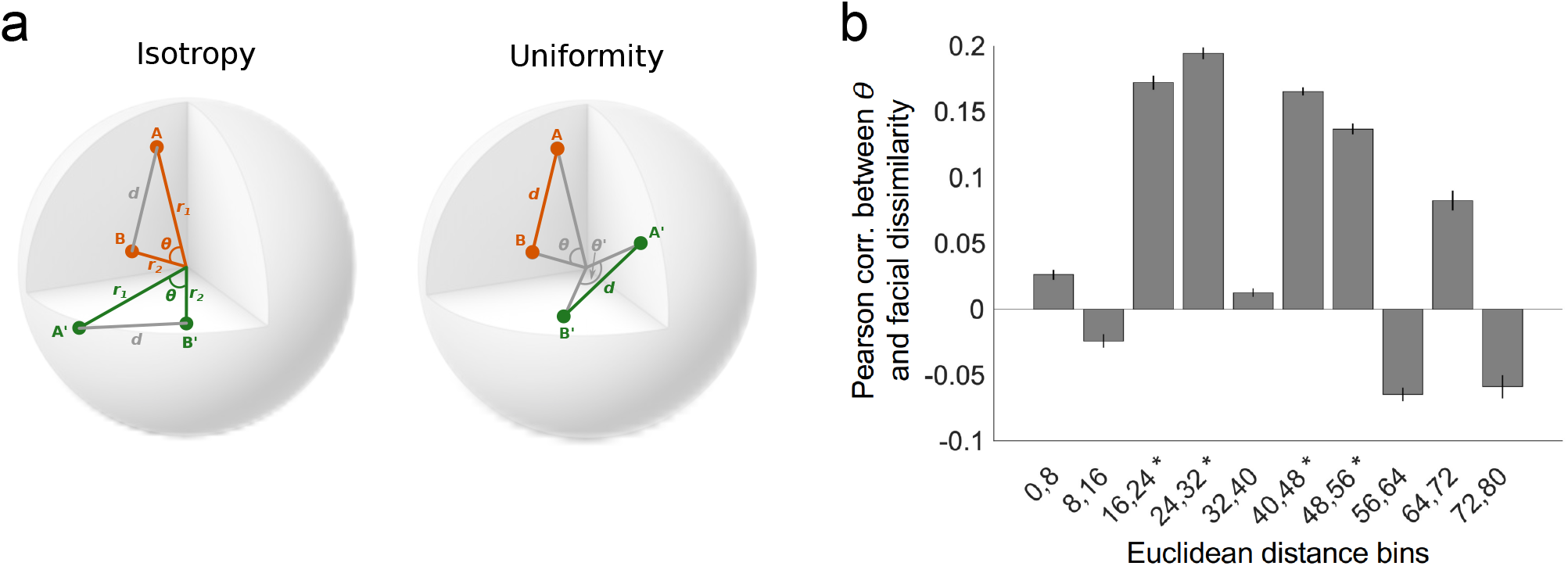
Uniformity test. **a)** Schematic of the difference between perceptual isotropy and perceptual uniformity. If space is perceptually isotropic (left), the pairs of faces (A,B) and (A’,B’) should appear equally dissimilar, because they correspond to vectors that have been rotated around the origin while preserving their geometric relationship to one another (the vectors span the same angle *θ* and have the same norms *r*_1_ and *r*_2_). If space is perceptually uniform (right), the pairs of faces A-B and A’-B’ should appear equally dissimilar, because they correspond to vectors that have been linearly translated in the space, preserving their Euclidean distance to one another (while disrupting their geometric relationship). **b)** Analysis evaluating evidence of perceptual uniformity in the stimulus set A experiment. Face pairs were binned into groups with similar Euclidean distances. We then evaluated whether the angle between the faces explains variance in perceived dissimilarity within each bin. If the space is non-uniform, we might expect faces with larger angular differences to appear more different, even if they have identical Euclidean distance. We find only weak evidence for any non-uniformity in the face space.

**Supplementary Fig. 3.**
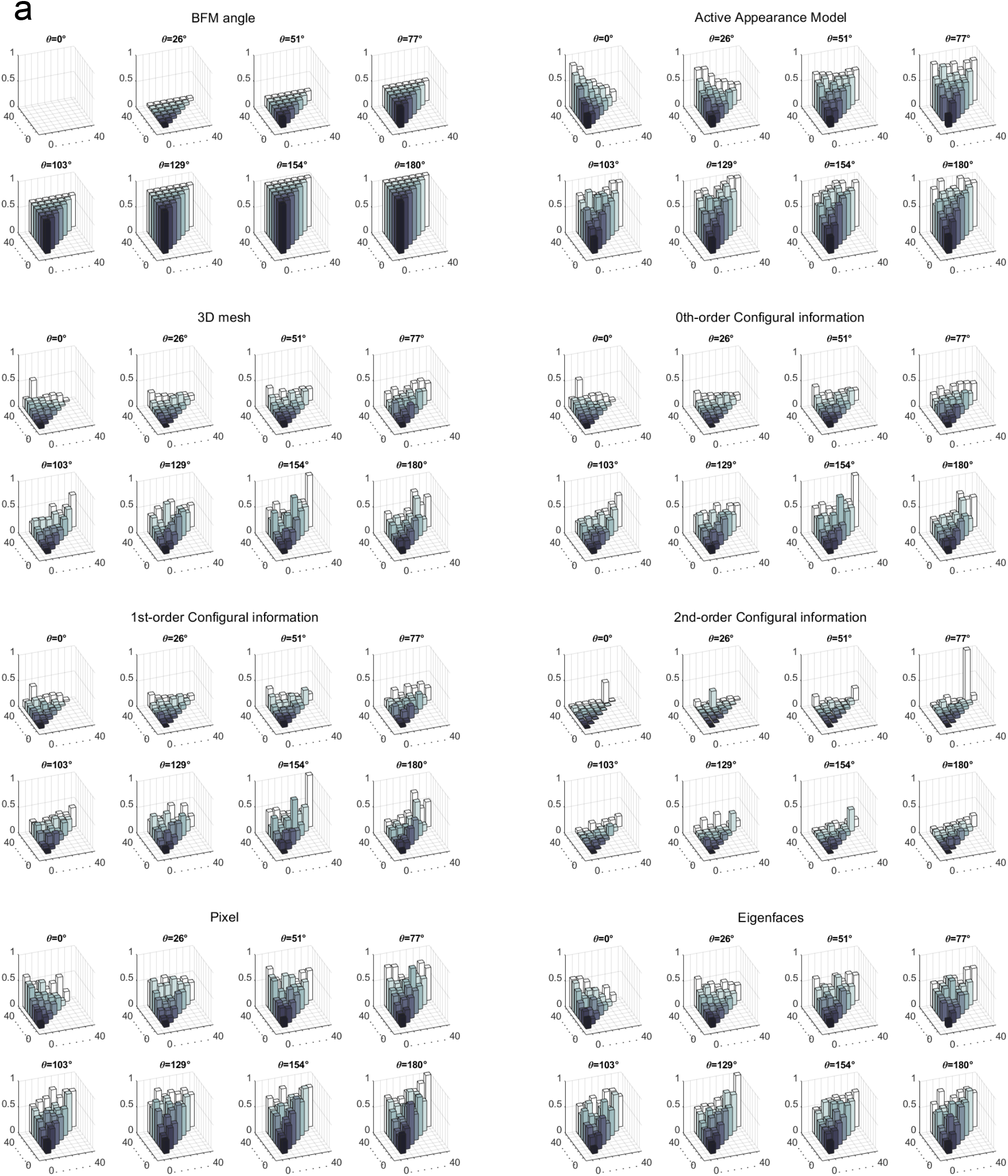
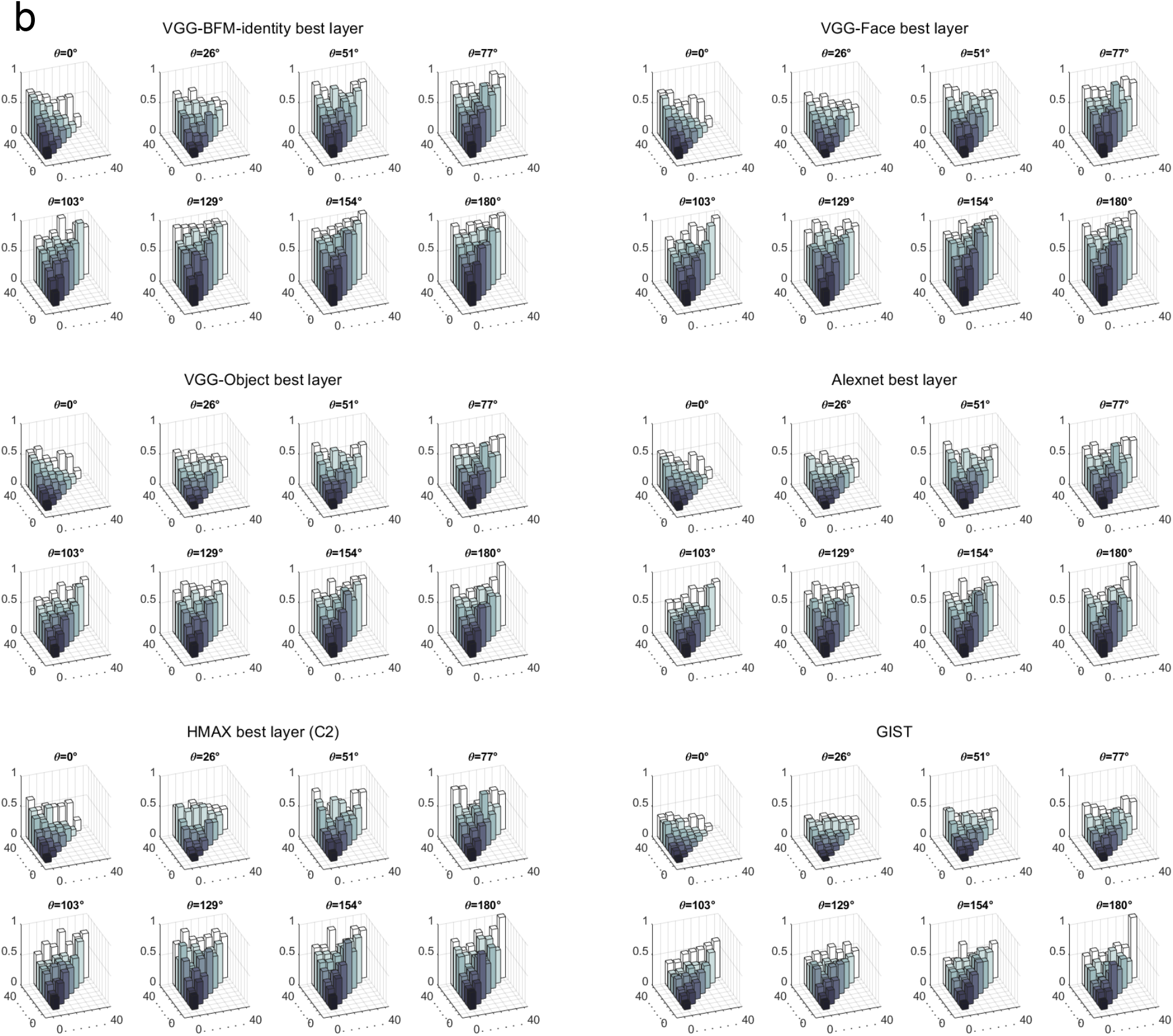
Dissimilarity predictions for each face pair in the stimulus set A as a function of angular and radial geometry in BFM, according to each model. **a)** Models based on BFM information, facial geometry, landmarks or configurations, and simple 2D image properties. **b)** Models based on deep neural network representations or shallower computer vision features. Conventions are as in Figure 2a of the main manuscript. Each plot shows a “slice” through the BFM face space, comprising stimuli separated by the same BFM angle. The x and y axes indicate the length of the longer and shorter radius in the face pair, and the height of the bar indicates predicted dissimilarity for the face pair according to the model, normalised to the range 0-1.

**Supplementary Fig. 4.**
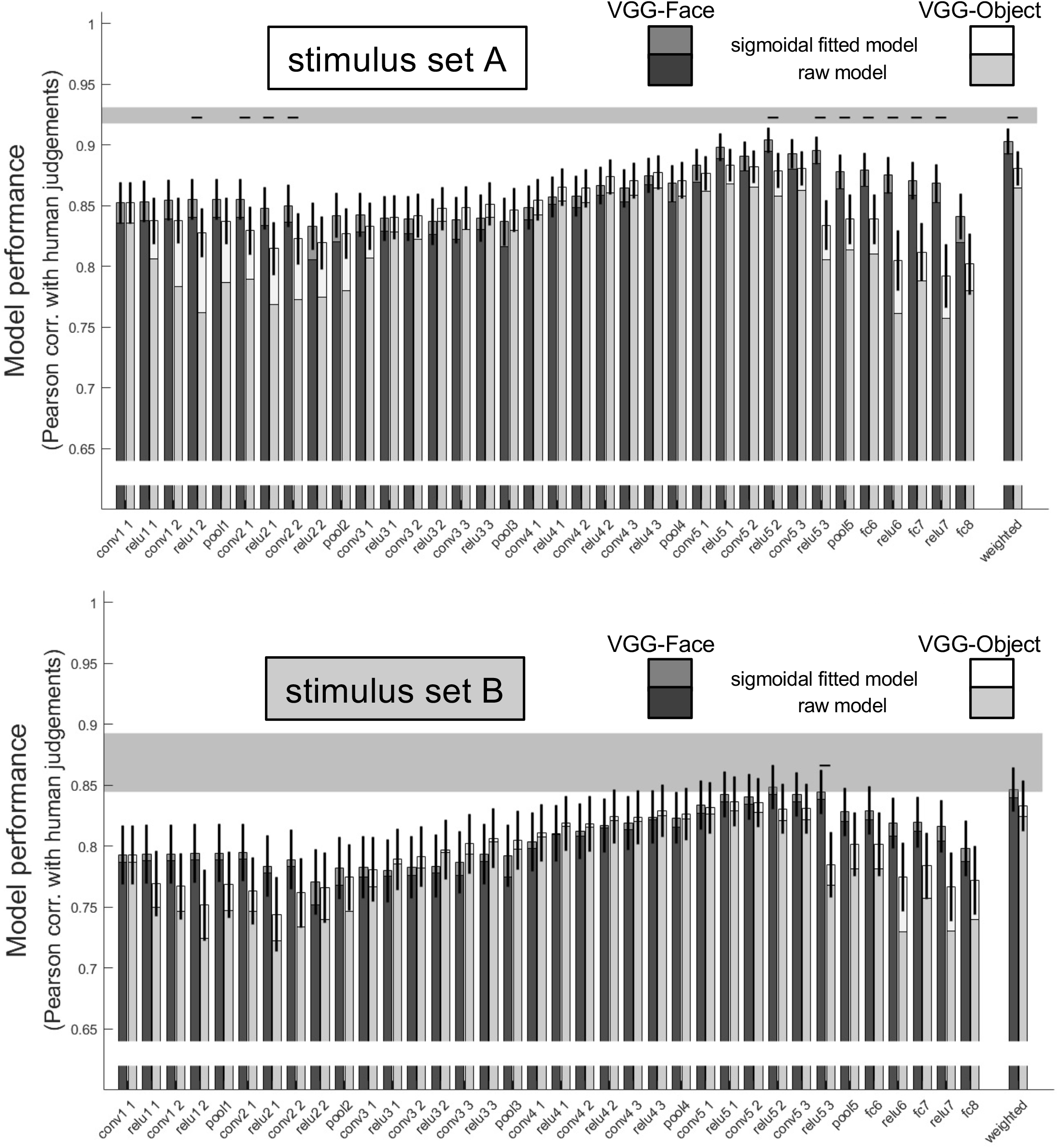
Ability of each layer within VGG-Face and VGG-Object neural networks to predict human face dissimilarity judgements. Correlations between human dissimilarity judgements in the stimulus set A (top) and B (bottom) and Euclidean distance within each layer of the same deep neural network trained either to recognise faces (VGG-Face) or objects (VGG-Object). All key processing steps within each network are shown, including application of a non-linearity (‘relu’), convolutional layers (‘conv’), max-pooling (‘pool’), and fully-connected layers (‘fc’). The darker lower part of each bar shows the performance of raw predicted distances, and paler upper parts show the same after fitting a sigmoidal transform, cross-validated over both participants and stimuli. The final bars show the performance of a linearly-weighted combination of all layers. Conventions are as in Figure 4b.

**Supplementary Fig. 5.**
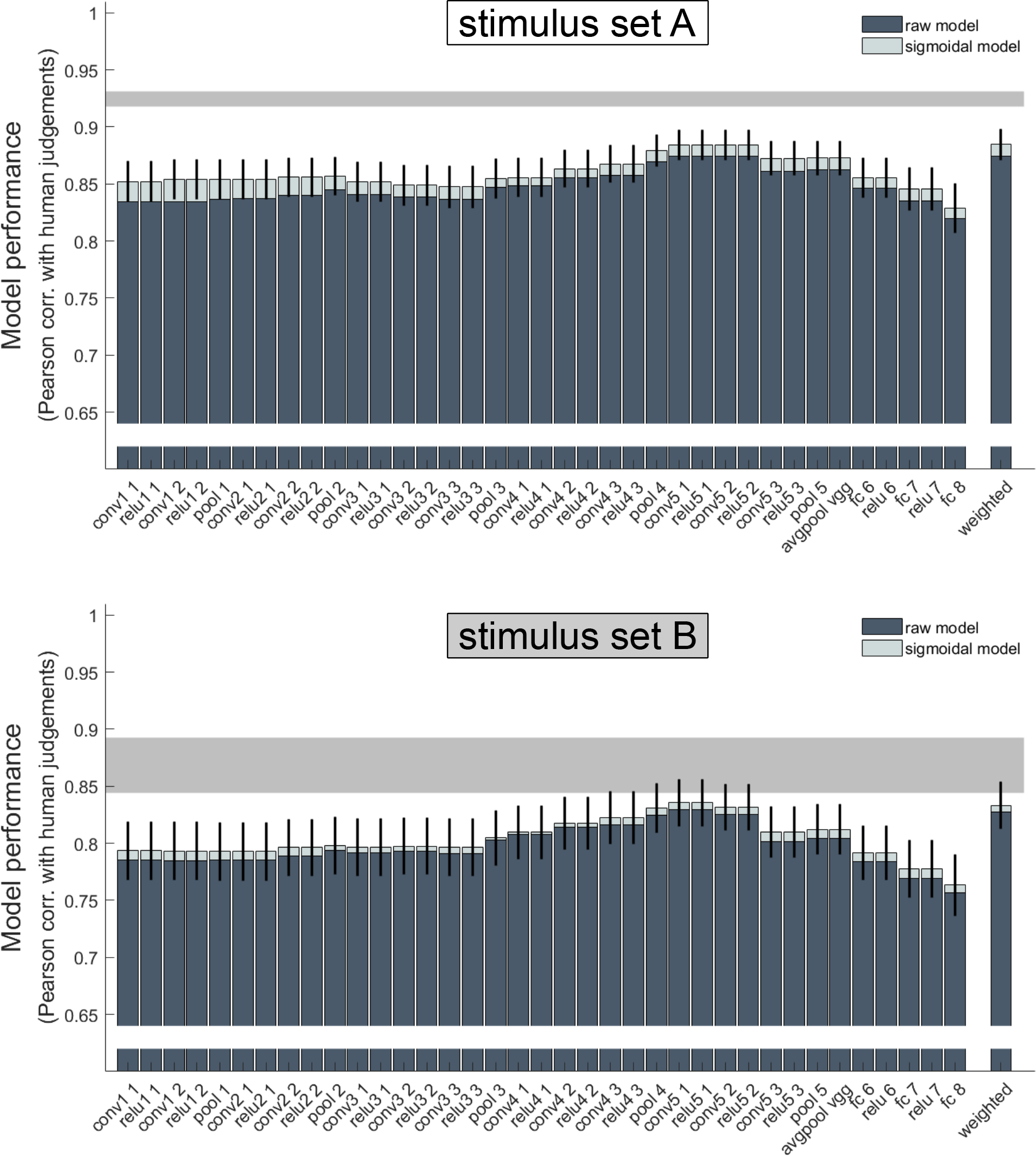
Ability of each layer within VGG-BFM-identity neural network to predict human face dissimilarity judgements. Correlations between human dissimilarity judgements in the stimulus set A (top) and B (bottom) and Euclidean distance within each layer of VGG-BFM-identity neural network. All key processing steps within each network are shown, including application of a non-linearity (‘relu’), convolutional layers (‘conv’), max-pooling (‘pool’), and fully-connected layers (‘fc’). The darker lower part of each bar shows the performance of raw predicted distances, and paler upper parts show the same after fitting a sigmoidal transform, cross-validated over both participants and stimuli. The final bars show the performance of a linearly-weighted combination of all layers. Conventions are as in Figure 4b.

**Supplementary Fig. 6.**
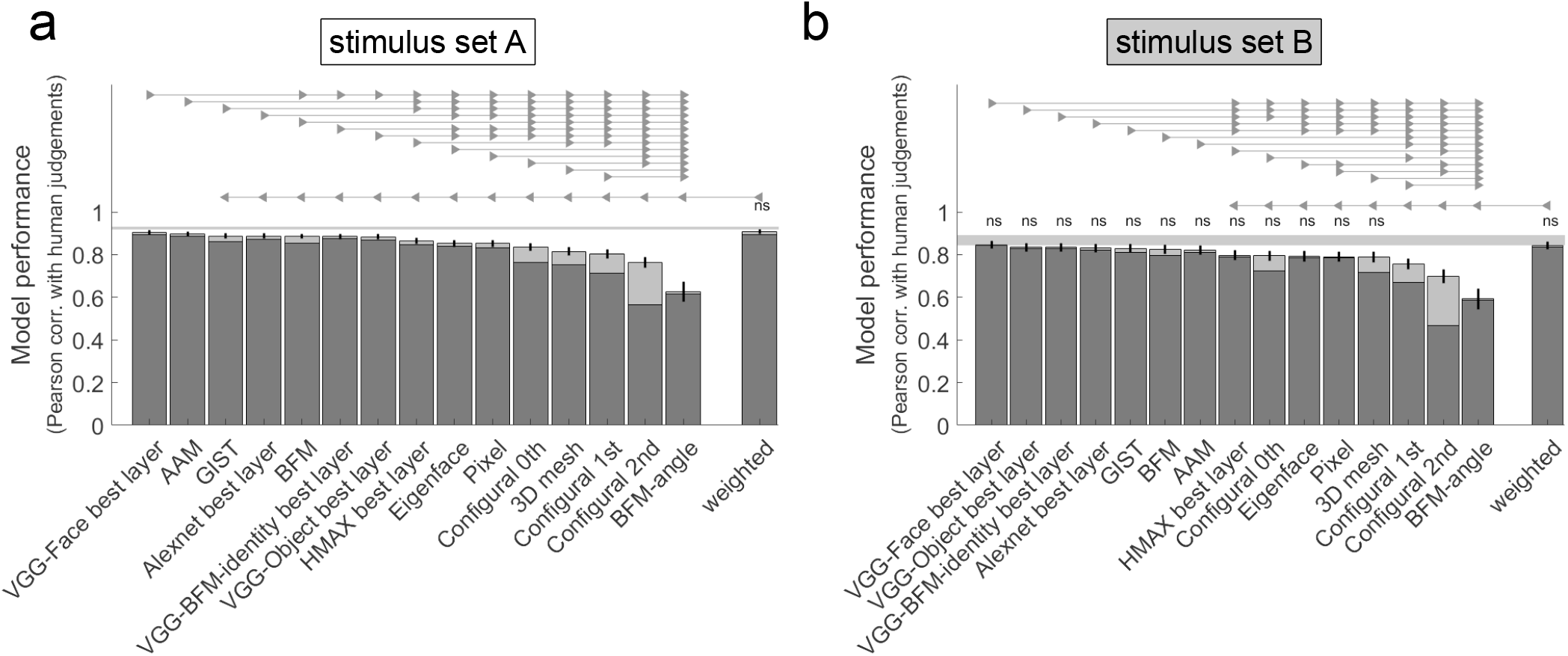
Statistical comparisons between models after allowing a sigmoidal transformation to fit human dissimilarity data. **a)** Model performance data shown in Figure 4b, but with models ordered and statistically compared according to their performance after fitting a sigmoidal transform (within cross-validation folds) to raw model-predicted distances. Conventions are as in Figure 4b. **b)** Corresponding data for stimulus set B.

**Supplementary Fig. 7.**
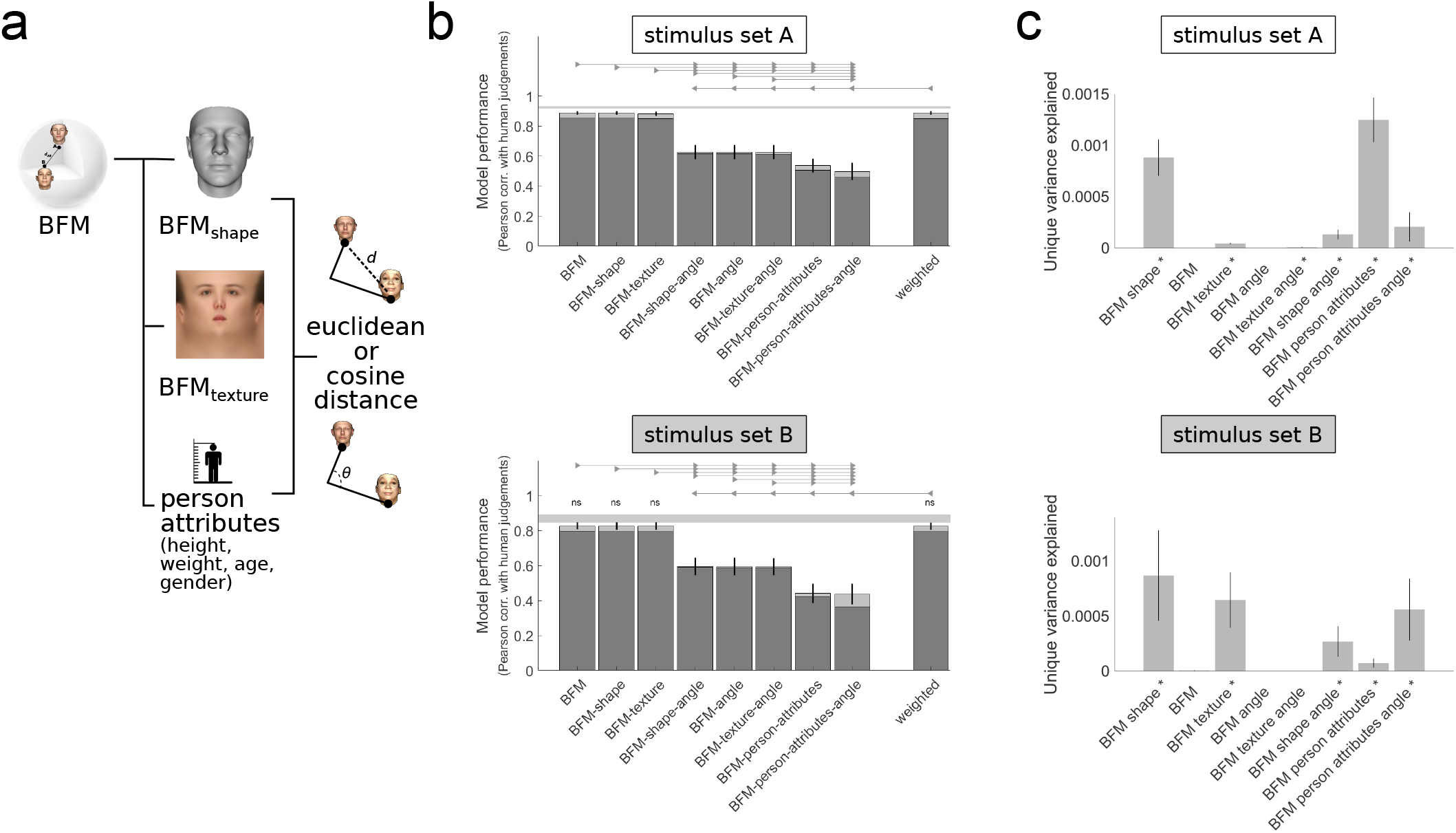
Performance of BFM models based on different subspaces and distance metrics. **a)** Schematic of the six additional models presented here and their relationship to the BFM models presented in the main manuscript. Here we evaluate again the performance of a model based on either Euclidean or Cosine distance in the full 398-dimensional BFM space, as well as ones based on Euclidean or Cosine distance in: (top) the 199-dimensional shape subspace, (middle) the 199-dimensional texture subspace, or (bottom) the 4-dimensional space comprising the dimensions that capture the most variance in height, weight, age, and gender. **b)** Average Pearson correlation of each model’s predictions with human data in the two experimental datasets. Conventions are as in Figure 4b. **c)** Unique variance analysis for the same models. Conventions are as in Figure 4c.

**Supplementary Fig. 8.**
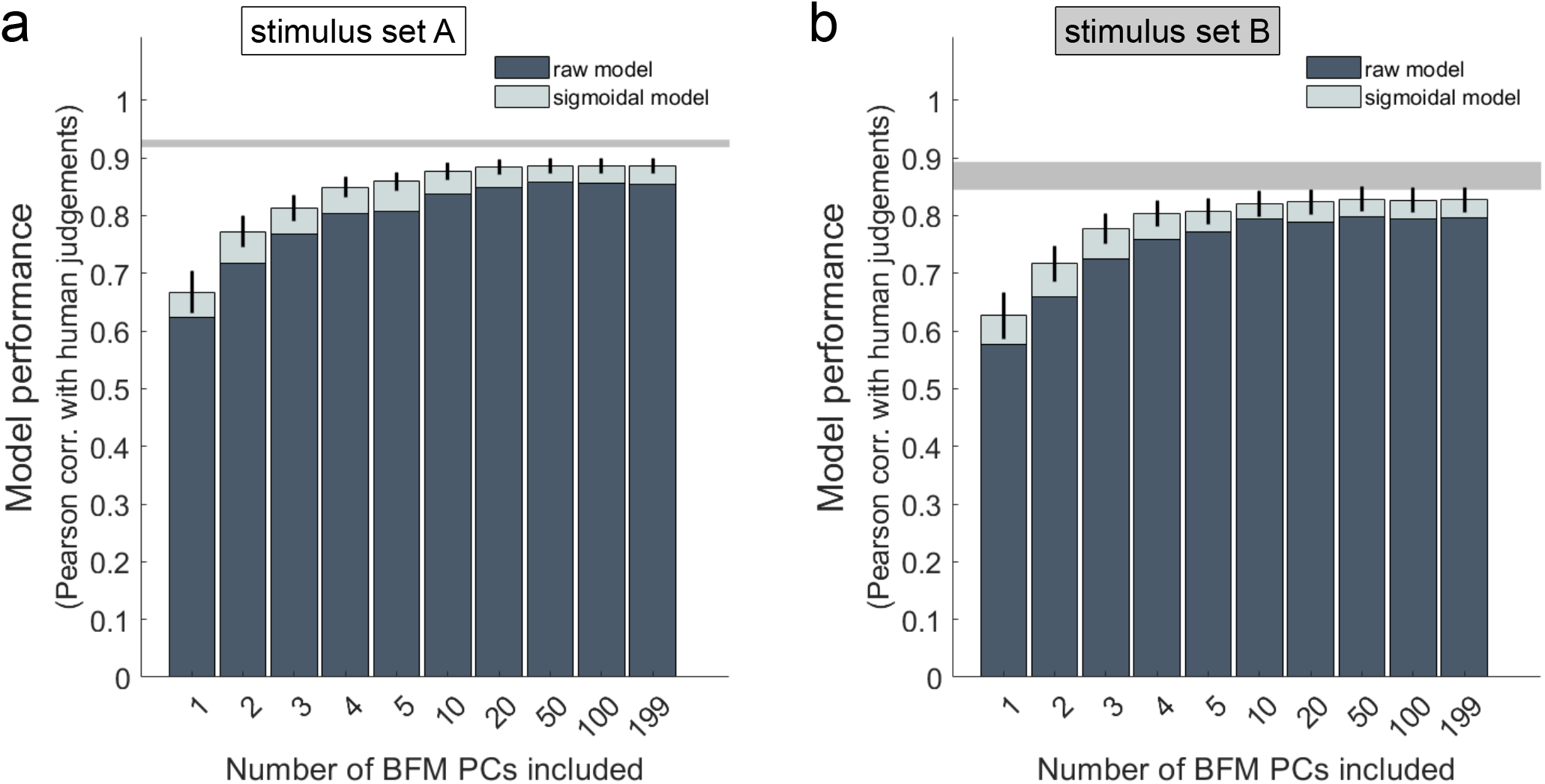
Effect of varying number of principal components in the 3D morphable Basel Face Model. **a)** Correlation with human dissimilarity ratings of the stimulus set A, for models comprising the first 1-5, 10, 20, 50, 100, or 199 principal components from both the shape and texture sub-spaces. For example, the 1-PC model measures the Euclidean distance between faces in a two-dimensional space consisting of the first shape-PC and first texture-PC. The 199-PC model is the full BFM space included in the main manuscript analyses. Including a larger number of components aids prediction, but performance rises rapidly and saturates at a smaller dimensionality than the full BFM space. **b)** Corresponding data for stimulus set B.

**Supplementary Fig. 9.**
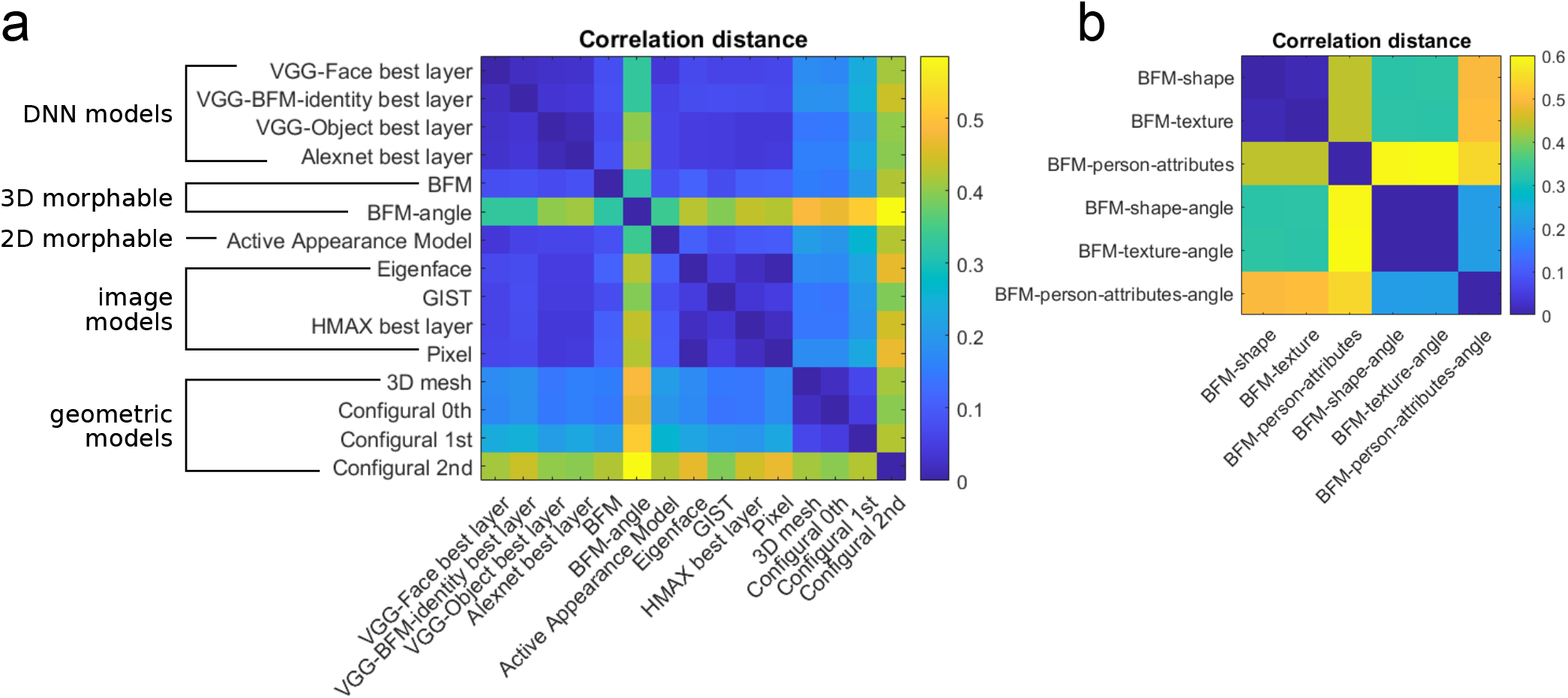
Correlations between model predictions for face pairs in Stimulus Set A. **a)** Models shown in the main manuscript (Figure 4), grouped by whether they derive from DNNs, from the principal components of the BFM, from simple image-computable models, or from the 3D geometry of faces. **b)** Models based on the BFM (see Supplementary Figure 7 for model performances).

## References

1. Nancy Kanwisher. Domain specificity in face perception. Nature neuroscience, 3(8):759–763, 2000.

2. Gillian Rhodes and David A Leopold. Adaptive norm-based coding of face identity. The Oxford handbook of face perception, pages 263–286, 2011.

3. Doris Y Tsao and Margaret S Livingstone. Mechanisms of face perception. Annu. Rev. Neurosci., 31:411–437, 2008.

4. Martha J Farah, Kevin D Wilson, Maxwell Drain, and James N Tanaka. What is” special” about face perception? Psychological review, 105(3):482, 1998.

5. Yann LeCun, Yoshua Bengio, and Geoffrey Hinton. Deep learning. nature, 521(7553):436–444, 2015.

6. Katherine R. Storrs and Nikolaus Kriegeskorte. Deep learning for cognitive neuroscience. In The Cognitive Neurosciences. 2020.

7. Olga Russakovsky, Jia Deng, Hao Su, Jonathan Krause, Sanjeev Satheesh, Sean Ma, Zhiheng Huang, Andrej Karpathy, Aditya Khosla, Michael Bernstein, et al. Imagenet large scale visual recognition challenge. International journal of computer vision, 115(3):211–252, 2015.

8. Pascal Paysan, Reinhard Knothe, Brian Amberg, Sami Romdhani, and Thomas Vetter. A 3d face model for pose and illumination invariant face recognition. In 2009 Sixth IEEE International Conference on Advanced Video and Signal Based Surveillance, pages 296–301. Ieee, 2009.

9. Bernhard Egger, William A. P. Smith, Ayush Tewari, Stefanie Wuhrer, Michael Zollhoefer, Thabo Beeler, Florian Bernard, Timo Bolkart, Adam Kortylewski, Sami Romdhani, Christian Theobalt, Volker Blanz, and Thomas Vetter. 3D Morphable Face Models—Past, Present, and Future. ACM Transactions on Graphics, 39(5):1–38, September 2020.

10. Ilker Yildirim, Mario Belledonne, Winrich Freiwald, and Josh Tenenbaum. Efficient inverse graphics in biological face processing. Science Advances, 6(10), 2020.

11. Tim Valentine. A unified account of the effects of distinctiveness, inversion, and race in face recognition. The Quarterly Journal of Experimental Psychology Section A, 43(2):161–204, 1991.

12. Tim Valentine, Michael B Lewis, and Peter J Hills. Face-space: A unifying concept in face recognition research. Quarterly Journal of Experimental Psychology, 69(10):1996–2019, 2016.

13. A Mike Burton and John R Vokey. The face-space typicality paradox: Understanding the face-space metaphor. The Quarterly Journal of Experimental Psychology Section A, 51(3):475–483, 1998.

14. Guy Wallis. Toward a unified model of face and object recognition in the human visual system. Frontiers in psychology, 4:497, 2013.

15. David A Leopold, Igor V Bondar, and Martin A Giese. Norm-based face encoding by single neurons in the monkey inferotemporal cortex. Nature, 442(7102):572–575, 2006.

16. David A Leopold, Alice J O’Toole, Thomas Vetter, and Volker Blanz. Prototype-referenced shape encoding revealed by high-level aftereffects. Nature neuroscience, 4(1):89–94, 2001.

17. Omkar M Parkhi, Andrea Vedaldi, and Andrew Zisserman. Deep face recognition. arXiv, 2015.

18. Jordan W Suchow, Joshua C Peterson, and Thomas L Griffiths. Learning a face space for experiments on human identity. 40th Annual Meeting of the Cognitive Science Society, 2018.

19. Matthew Q. Hill, Connor J. Parde, Carlos D. Castillo, Y. Ivette Colón, Rajeev Ranjan, Jun-Cheng Chen, Volker Blanz, and Alice J. O’Toole. Deep convolutional neural networks in the face of caricature. Nature Machine Intelligence, 1(11):522–529, November 2019. ISSN 2522-5839. doi: 10.1038/s42256-019-0111-7.

20. Nadine Gummersbach and Volker Blanz. A morphing-based analysis of the perceptual distance metric of human faces. Proceedings of the 7th Symposium on Applied Perception in Graphics and Visualization, page 109, 2010.

21. Christoph Daube, Tian Xu, Jiayu Zhan, Andrew Webb, Robin A.A. Ince, Oliver G.B. Garrod, and Philippe G. Schyns. Grounding deep neural network predictions of human categorization behavior in understandable functional features: The case of face identity. Patterns, page 100348, September 2021.

22. Johan D. Carlin and Nikolaus Kriegeskorte. Adjudicating between face-coding models with individual-face fMRI responses. PLOS Computational Biology, 13(7):e1005604, 2017.

23. Neil MT Houlsby, Ferenc Huszár, Mohammad M Ghassemi, Gergo? Orbán, Daniel M Wolpert, and Máté Lengyel. Cognitive tomography reveals complex, task-independent mental representations. Current Biology, 23(21):2169–2175, 2013.

24. Chaitanya K Ryali, Xiaotian Wang, and Angela J Yu. Leveraging Computer Vision Face Representation to Understand Human Face Representation. Cogsci., page 15, 2021.

25. Timothy F. Cootes, Gareth J. Edwards, and Christopher J. Taylor. Active appearance models. IEEE Transactions on pattern analysis and machine intelligence, 23(6):681–685, 2001.

26. Joan Alabort-i Medina and Stefanos Zafeiriou. A unified framework for compositional fitting of active appearance models. International Journal of Computer Vision, 121(1):26–64, 2017.

27. Marieke Mur, Mirjam Meys, Jerzy Bodurka, Rainer Goebel, Peter a. Bandettini, and Nikolaus Kriegeskorte. Human object-similarity judgments reflect and transcend the primate-IT object representation. Frontiers in Psychology, 4:1–22, 2013.

28. Karen Simonyan and Andrew Zisserman. Very deep convolutional networks for large-scale image recognition. arXiv, 2014.

29. Alex Krizhevsky, Ilya Sutskever, and Geoffrey E Hinton. Imagenet classification with deep convolutional neural networks. In Advances in neural information processing systems, pages 1097–1105, 2012.

30. Le Chang and Doris Y. Tsao. The Code for Facial Identity in the Primate Brain. Cell, 169(6):1013–1028.e14, 2017.

31. Pawan Sinha, Benjamin Balas, Yuri Ostrovsky, and Richard Russell. Face recognition by humans: Nineteen results all computer vision researchers should know about. Proceedings of the IEEE, 94 (11):1948–1962, 2006.

32. Gillian Rhodes. Looking at faces: First-order and second-order features as determinants of facial appearance. Perception, 17(1):43–63, 1988.

33. Steven C Dakin and Roger J Watt. Biological “bar codes” in human faces. Journal of vision, 9(4):2–2, 2009.

34. Aleix M. Martinez and Shichuan Du. A model of the perception of facial expressions of emotion by humans: Research overview and perspectives. Journal of Machine Learning Research, 2017.

35. Irving Biederman and Peter Kalocsais. Neurocomputational bases of object and face recognition. Philosophical Transactions of the Royal Society of London. Series B: Biological Sciences, 352 (1358):1203–1219, 1997.

36. Xiaomin Yue, Irving Biederman, Michael C Mangini, Christoph von der Malsburg, and Ori Amir. Predicting the psychophysical similarity of faces and non-face complex shapes by image-based measures. Vision research, 55:41–46, 2012.

37. Mark Steyvers and Tom Busey. Predicting similarity ratings to faces using physical descriptions. Computational, geometric, and process perspectives on facial cognition: Contexts and challenges, pages 115–146, 2000.

38. Tal Golan, Prashant C. Raju, and Nikolaus Kriegeskorte. Controversial stimuli: Pitting neural networks against each other as models of human cognition. Proceedings of the National Academy of Sciences, 117(47):29330–29337, 2020.

39. Ryan M. Stolier, Eric Hehman, Matthias D. Keller, Mirella Walker, and Jonathan B. Freeman. The conceptual structure of face impressions. Proceedings of the National Academy of Sciences, 115 (37):9210–9215, 2018.

40. Selma Carolin Rudert, Leonie Reutner, Rainer Greifeneder, and Mirella Walker. Faced with exclusion: Perceived facial warmth and competence influence moral judgments of social exclusion. Journal of Experimental Social Psychology, 68:101–112, 2017.

41. Rainer Scheuchenpflug. Predicting face similarity judgements with a computational model of face space. Acta psychologica, 100(3):229–242, 1999.

42. Alice J O’Toole, Carlos D Castillo, Connor J Parde, Matthew Q Hill, and Rama Chellappa. Face space representations in deep convolutional neural networks. Trends in cognitive sciences, 22(9): 794–809, 2018.

43. Kieran Lee, Graham Byatt, and Gillian Rhodes. Caricature effects, distinctiveness, and identification: Testing the face-space framework. Psychological science, 11(5):379–385, 2000.

44. Volker Blanz, Alice J O’toole, Thomas Vetter, and Heather A Wild. On the other side of the mean: The perception of dissimilarity in human faces. Perception, 29(8):885–891, 2000.

45. Fang Jiang, Volker Blanz, and Alice J O’Toole. Probing the visual representation of faces with adaptation: A view from the other side of the mean. Psychological Science, 17(6):493–500, 2006.

46. David A Ross, Mickael Deroche, and Thomas J Palmeri. Not just the norm: Exemplar-based models also predict face aftereffects. Psychonomic bulletin & review, 21(1):47–70, 2014.

47. Katherine R Storrs and Derek H Arnold. Not all face aftereffects are equal. Vision research, 64:7–16, 2012.

48. H Hill. Information and viewpoint dependence in face recognition. Cognition, 62(2):201–222, 1997.

49. Chang Hong Liu, Charles A Collin, A. Mike Burton, and Avi Chaudhuri. Lighting direction affects recognition of untextured faces in photographic positive and negative. Vision Research, 39(24):4003–4009, 1999.

50. Vicki Bruce. Changing faces: Visual and non-visual coding processes in face recognition. British Journal of Psychology, 73(1):105–116, 1982.

51. James J DiCarlo and David D Cox. Untangling invariant object recognition. Trends in cognitive sciences, 11(8):333–341, 2007.

52. Katharina Dobs, Leyla Isik, Dimitrios Pantazis, and Nancy Kanwisher. How face perception unfolds over time. Nature Communications, 10(1):1258, 2019.

53. Kamila Maria Jozwik, Martin Schrimpf, Nancy Kanwisher, and James J. DiCarlo. To find better neural network models of human vision, find better neural network models of primate vision. bioRxiv, 2019.

54. Kamila Maria Jozwik, Michael Lee, Tiago Marques, Martin Schrimpf, and Pouya Bashivan. Large-scale hyperparameter search for predicting human brain responses in the Algonauts challenge. bioRxiv, 2019.

55. John C Brigham and Roy S Malpass. The role of experience and contact in the recognition of faces of own-and other-race persons. Journal of social issues, 41(3):139–155, 1985.

56. Robert K Bothwell, John C Brigham, and Roy S Malpass. Cross-racial identification. Personality and Social Psychology Bulletin, 15(1):19–25, 1989.

57. Zhou Wang and Eero P Simoncelli. Maximum differentiation (mad) competition: A methodology for comparing computational models of perceptual quantities. Journal of Vision, 8(12):8–8, 2008.

58. Kamila Maria Jozwik, Ian Charest, Nikolaus Kriegeskorte, and Radoslaw Martin Cichy. Animacy dimensions ratings and approach for decorrelating stimuli dimensions. bioRxiv, 2021.

59. Hamed Nili, Cai Wingfield, Alexander Walther, Li Su, William Marslen-Wilson, and Nikolaus Kriegeskorte. A toolbox for representational similarity analysis. PLoS Computational Biology, 10(4), 2014.

60. Katherine R. Storrs, Tim C. Kietzmann, Alexander Walther, Johannes Mehrer, and Nikolaus Kriegeskorte. Diverse Deep Neural Networks All Predict Human Inferior Temporal Cortex Well, After Training and Fitting. Journal of Cognitive Neuroscience, 33(10), 2021.

61. Katherine R Storrs, Seyed-Mahdi Khaligh-Razavi, and Nikolaus Kriegeskorte. Noise ceiling on the crossvalidated performance of reweighted models of representational dissimilarity: Addendum to khaligh-razavi & kriegeskorte (2014). bioRxiv, 2020.

62. Tim C Kietzmann, Courtney J Spoerer, Lynn Sörensen, Radoslaw M Cichy, Olaf Hauk, and Nikolaus Kriegeskorte. Recurrence required to capture the dynamic computations of the human ventral visual stream. Proceedings of the National Academy of Sciences, 2019.

63. Ziwei Liu, Ping Luo, Xiaogang Wang, and Xiaoou Tang. Deep learning face attributes in the wild. Proceedings of International Conference on Computer Vision (ICCV), December 2015.

64. Maximilian Riesenhuber and Tomaso Poggio. Hierarchical models of object recognition in cortex. Nature neuroscience, 2(11):1019–1025, 1999.

65. Aude Oliva and Antonio Torralba. Modeling the shape of the scene: A holistic representation of the spatial envelope. International journal of computer vision, 42(3):145–175, 2001.

66. Jessica Taubert, Deborah Apthorp, David Aagten-Murphy, and David Alais. The role of holistic processing in face perception: Evidence from the face inversion effect. Vision research, 51(11): 1273–1278, 2011.

67. Alexander Buslaev, Alex Parinov, Eugene Khvedchenya, Vladimir I. Iglovikov, and Alexandr A. Kalinin. Albumentations: fast and flexible image augmentations. Information, 11(2):125, 2020.

